# Advances and limits of species-level canopy mapping in a hyperdiverse tropical forest: multi-temporal crown segmentation and airborne imaging spectroscopy

**DOI:** 10.1101/2024.06.24.600405

**Authors:** James G C Ball, Sadiq Jaffer, Anthony Laybros, Colin Prieur, Toby Jackson, Anil Madhavapeddy, Nicolas Barbier, David A Coomes, Grégoire Vincent

**Author notes:** Corresponding author(s)*: James G C Ball; Grégoire Vincent.

## Abstract

Species-level canopy maps underpin tropical forest biodiversity monitoring, conservation planning, and carbon accounting, yet high species richness and structural complexity make remote classification challenging. Here we evaluate how far a two-step mapping approach can be taken, and where it breaks down, in hyperdiverse moist forest at the Paracou Field Station, French Guiana. First, we delineate individual tree crowns from ten repeat uncrewed aerial vehicle (UAV) RGB surveys with Mask R-CNN, fusing predictions across dates by temporal consensus: mean segmentation F1 rose from 0.68 (single date) to 0.78 (ten dates), covering approximately 86% of test-region canopy area. Second, we classify each crown from a single airborne hyperspectral acquisition (416–2500 nm, 1 m) using machine learning classifiers trained and tested on 3,186 field-verified crowns spanning 169 species. Linear Discriminant Analysis performed best (weighted F1 = 0.75), outperforming more flexible models, but unevenly: across repeated cross-validation (20 × 5-fold), on average 50 species (95% CI: 41–63) attained F1 ≥ 0.7 in a given fold and only 15 did so reliably, with many rare species unclassifiable (macro-average F1 = 0.48). Combining both steps, we estimate approximately 70% of the landscape’s canopy area was correctly mapped to species. Band-importance and ablation analyses identified the far-red edge (748–775 nm) as the most informative spectral region. These results advance on studies limited to 20 or fewer species and set out spectral-resolution and training-data requirements for airborne and forthcoming spaceborne imaging spectrometers, while showing that accuracy remains strongly conditioned by training-data availability and the single-site, single-acquisition design.

## 1 Introduction

Landscape-scale maps of forest species have many applications in ecology, forestry, and conservation [Chave et al., 2003, Fassnacht et al., 2016, Davies et al., 2021, Weinstein et al., 2024]. By resolving where individual species occur, they enable tracking of species abundance and health [Baldeck et al., 2015, Park et al., 2019], support conservation of threatened taxa [McGrath et al., 2009, Tuomisto et al., 2003], and aid monitoring of invasive species and disease spread [Chance et al., 2016, Liu et al., 2021, Chan et al., 2021]. Species-resolved inventories also underpin sustainable forestry [Laybros et al., 2020, White et al., 2016], improve carbon-stock estimates [Slik et al., 2013], support forest type classification [Vaglio Laurin et al., 2014, Bohlman, 2015], and reveal links between biodiversity and ecosystem functioning [Reichstein et al., 2013, Kamoske et al., 2022], including individual-level phenology and productivity [Park et al., 2019, Schmitt et al., 2024]. Species-level spatial data also remain central for understanding biodiversity maintenance through density-dependent mortality, dispersal limitation, and succession [Kenkel, 1988, LaManna et al., 2017, Barry and Schnitzer, 2021], and for tracking physiological responses and forest resilience [e.g., Nepstad et al., 2007, Anderegg et al., 2015, Rowland et al., 2015]. However, existing methods have been demonstrated mainly at local scales and under favourable conditions (gentle terrain, low species mixture, few age classes; Fassnacht et al. 2016). Hyperdiverse moist tropical forests, with their complex canopy structure and high species richness, remain a frontier for operational species mapping.

Species-level mapping typically involves two steps: *individual tree crown (ITC) delineation* (or segmentation), which partitions the canopy into discrete crown polygons, and *species classification*, which assigns a species identity to each crown. Pixel- or patch-based approaches can bypass explicit ITC delineation [Tang et al., 2021], but crown-based workflows remain standard for tree-level ecological inference. Segmentation draws on lidar, imagery, or both, while species classification typically relies on hyperspectral remote sensing. Obstacles to accurate mapping persist at both stages, but are most prominent during the latter.

For segmentation, airborne lidar methods are effective in temperate and boreal forests [e.g., Dalponte and Coomes, 2016, Hastings et al., 2020] but struggle in complex tropical canopies where crowns interweave and subcanopy trees are largely invisible [Aubry-Kientz et al., 2019, Cao et al., 2023]. Image-based methods use high-resolution RGB colour and texture, often with neural networks such as Mask R-CNN [He et al., 2017], to separate irregular crown boundaries [Ball et al., 2023, Gan et al., 2023], but variable illumination, tree sway, phenological change, and ortho-mosaicking artefacts still reduce accuracy [Park et al., 2019, Gan et al., 2023].

One promising strategy for improving accuracy involves segmenting trees in imagery collected from multiple surveys conducted over time, then evaluating which crowns are consistently identified across these datasets [Nuijten et al., 2019]. This approach helps distinguish genuine tree crowns from artefacts caused by environmental changes or processing variability [Takahashi Miyoshi et al., 2020], but has yet to be critically evaluated.

For species identification, imaging spectrometers record canopy reflectance across many narrow bands, and species can be identified by their characteristic spectra, shaped by biochemical and structural traits [Ustin et al., 2004, Ollinger, 2011, Meireles et al., 2020]. This approach is most effective in low-diversity systems with limited spectral overlap among classes [Fassnacht et al., 2016]; high-diversity systems pose additional challenges. Canopy reflectance integrates multiple interacting traits [Curran, 1989, Jacquemoud and Baret, 1990], and these relationships differ across environments and canopy layers, making identification of the most informative spectral regions non-trivial [Schweiger et al., 2021]. Trait differences (pigments, structural compounds, water content) are often phylogenetically conserved, limiting separability among congeners [Cavender-Bares et al., 2016, Meireles et al., 2020], while environmental influences, phenology, and canopy-scale structure further modulate spectral expression [Nunes et al., 2017, Knyazikhin et al., 2013, Béland and Kobayashi, 2024]. The red edge and adjacent NIR region appear particularly important for species separation in diverse forests [Durgante et al., 2013, Hennessy et al., 2020, Pereira Martins-Neto et al., 2023], but the universality of this finding requires further study.

In practice, early studies using AVIRIS data have successfully classified 7 species in Costa Rica [Clark et al., 2005] and 17 in Hawaii [Féret and Asner, 2013]. More recent work in French Guiana classified 20 common species with ∼80% accuracy [Laybros et al., 2019], however this is still only a fraction of the species present in wet tropical forests (upwards of 100 species per hectare; Valencia et al. 1994, ter Steege et al. 2013). Advanced machine learning approaches – including convolutional neural networks applied to spectral profiles [Fricker et al., 2019], multi-layer perceptrons [Mäyrä et al., 2021], and more recently transformer architectures [Hong et al., 2024] – show promise given their capacity to learn non-linear spectral representations. However, their effectiveness relative to classical methods remains contested, particularly in high-diversity settings where training data per species are limited and overfitting is a concern [Féret and Asner, 2013].

Despite this growing body of work, important needs remain: multi-temporal imagery is rarely exploited to make crown delineation more robust in complex tropical canopies, classification in the wet tropics has largely been confined to 20 or fewer common species, and the spectral regions most critical for discrimination at this taxonomic scale have not been systematically validated through ablation.

Here we evaluate a two-step approach for species-level mapping at the Paracou Field Station, French Guiana, trained and tested on 3,186 manually delineated and field-verified crowns spanning 169 species (drawn from a labelled pool of 3,256 crowns across 239 species; see Section 2.7). We first delineated individual tree crowns from multi-temporal UAV RGB imagery using *detectree2*, a Mask R-CNN-based tool [He et al., 2017, Ball et al., 2023], and developed a consensus-fusion method that spatially matches crown polygons across dates and averages their vertices, exploiting the feasibility of repeat low-cost UAV surveys to address temporal variability in canopy appearance [Martin et al., 2018, Shamaoma et al., 2023]. We then classified the species of each crown from airborne hyperspectral data, comparing several classifiers and performing ablation experiments to identify the most informative spectral regions.

The research aims to:

1. Assess whether the accuracy of tree crown maps can be improved by making repeated aerial surveys, segmenting tree crowns in each survey, and combining segmentation results to construct a consensus-based map;
2. Evaluate which machine learning classifier yields the most accurate overall (landscape) predictions of the species identity of individual tree crowns and whether the inclusion of rarer species in training is detrimental to overall accuracy;
3. Determine which spectral regions are most important for species discrimination and validate these findings through ablation experiments.

The contribution is an end-to-end evaluation of how far current methods can take species-level mapping in a hyper-diverse tropical forest, anchored by an unusually large and carefully curated field dataset, quantifying the successes and the limitations of each step. A companion paper [Ball et al., 2026] examines the ecological and evolutionary determinants of species-level spectral separability using the same dataset.

## 2 Materials and Methods

### 2.1 Study site

The research was conducted at Paracou Field Station, French Guiana (5*^◦^*16’N 52*^◦^*55’W; see Fig. 1), where low-land tropical rainforest grows mostly on shallow ferralitic soils underlain by a variably transformed loamy saprolite [Gourlet-Fleury et al., 2004]. Mean annual rainfall is approximately 3200 mm, with a three-month dry season from mid-August to mid-November when rainfall is typically below 50 mm per month [Bonal et al., 2008, Wagner et al., 2011]. The station has 27 permanent plots of 0.5–25 ha containing approximately 76,000 trees of diameter at breast height (DBH) ≥ 10 cm and over 800 species [Gourlet-Fleury et al., 2004], inventoried every 1–5 years with the species, precise location and DBH of each trunk recorded. The ten most common species account for just over 30% of individuals, and 90% of species have been placed in a time-calibrated phylogeny by Baraloto et al. [2012].

**Figure 1:**
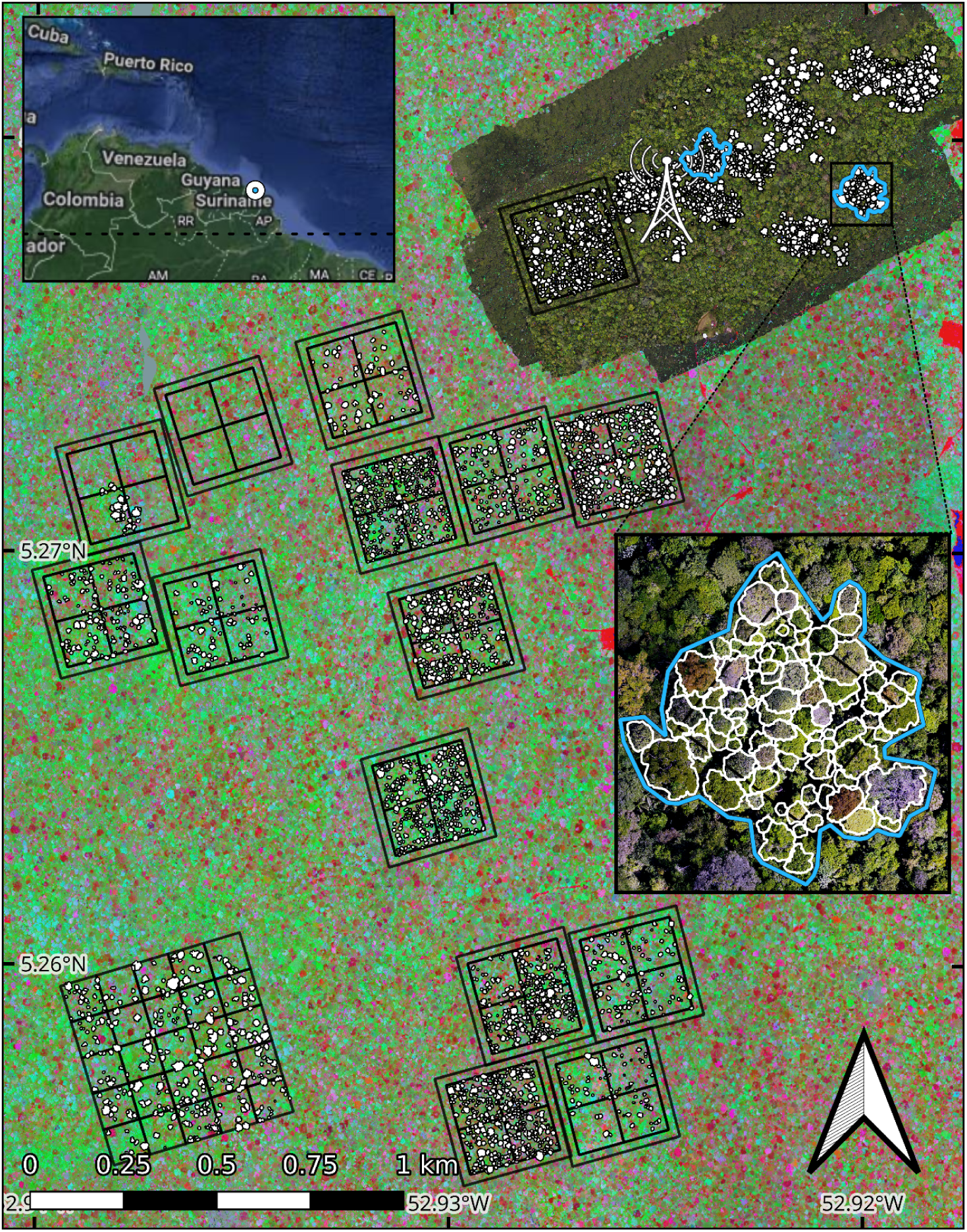
Site map of Paracou, French Guiana, with crowns and hyperspectral imagery. The manually delineated, labelled crowns are in white. The colourful background scan that covers the entire site is a representation of the hyperspectral data (selected projected PCA bands for illustration). The repeat survey UAV-RGB region is shown in the northwest around the site’s flux tower. Within this region the segmentation test data areas are delineated in blue - the crowns within these areas were excluded from all training of the segmentation delineation. One of these regions is shown up close in the inset. The black boxes show the plots in which inventories are conducted.

### 2.2 Overview of methods

We combined UAV RGB imagery and airborne imaging spectroscopy, both co-registered to a lidar-derived Canopy Height Model (CHM; Table 1, Fig. 2). A convolutional neural network (CNN) applied to 10 UAV surveys over 6 months located and delineated crowns, and hyperspectral imagery was used to classify their species from the spectral ‘signature’ each species produces across the 416–2500 nm range. Predictions were evaluated against unseen test sets of manual crowns never exposed to the algorithms during training.

**Figure 2:**
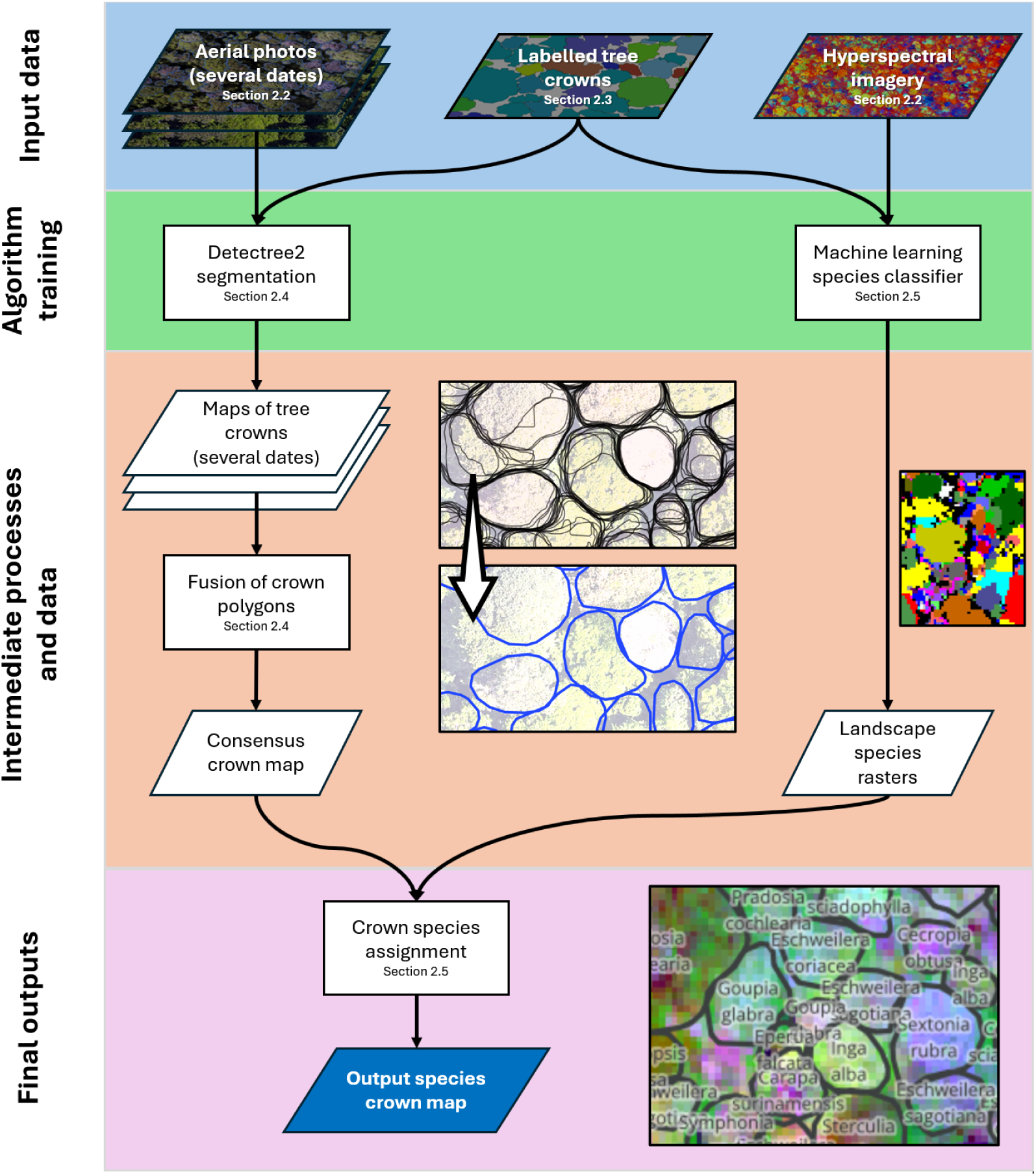
Simplified schematic of the crown mapping approach showing input data and the intermediate steps to producing a labelled tree crown map. In the centre is an illustration of the process of temporal polygon fusion. The top image shows the overlapping tree crown polygons predicted over multiple dates. Where polygons have a high degree of overlap, each will still have a slightly different shape due to differences in the RGB orthomosaics through time. The bottom image depicts the polygons after the fusion process, effectively averaging the positions of the vertices of the original polygons and discarding those without good consensus through time.

**Table 1:**
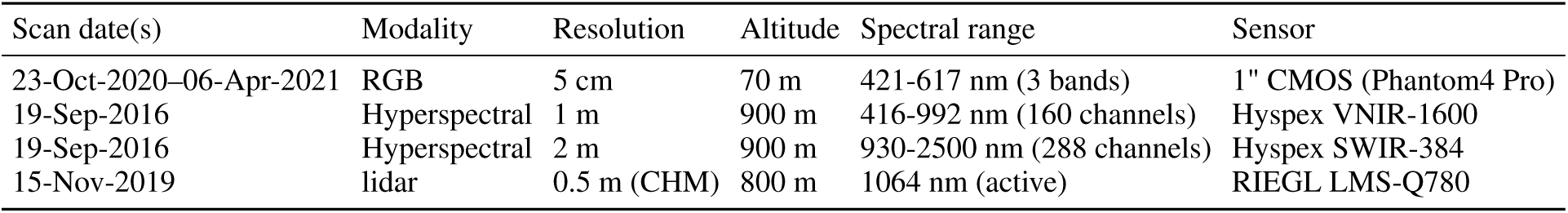
Primary remote sensing data sources used in the study (see Section 2.1 for the study location). Resolution is given as ground resolution for the RGB orthomosaic and as the processed CHM resolution for the lidar scans. Altitude is given as height above the forest canopy. An earlier lidar acquisition, contemporaneous with the 2016 hyperspectral campaign, was used solely to generate the surface model for hyperspectral orthorectification and is described in Section S2.

| Scan date(s) | Modality | Resolution | Altitude | Spectral range | Sensor |
| --- | --- | --- | --- | --- | --- |
| 23-Oct-2020–06-Apr-2021 | RGB | 5 cm | 70 m | 421–617 nm (3 bands) | 1" CMOS (Phantom4 Pro) |
| 19-Sep-2016 | Hyperspectral | 1 m | 900 m | 416–992 nm (160 channels) | Hypex VNIR-1600 |
| 19-Sep-2016 | Hyperspectral | 2 m | 900 m | 930–2500 nm (288 channels) | Hypex SWIR-384 |
| 15-Nov-2019 | lidar | 0.5 m (CHM) | 800 m | 1064 nm (active) | RIEGL LMS-Q780 |

### 2.3 Remote sensing data acquisition and co-registration

UAVs (DJI Phantom 4 Advanced and DJI P4 Multispectral) were employed to collect high-resolution RGB imagery, with a scan approximately every three weeks over a six-month period (10 surveys in total) of the region shown in Fig. 1. The RGB orthomosaics were compiled from the raw geotagged UAV photographs using structure from motion (SfM) photogrammetry in Agisoft Metashape. To improve spatio-temporal coherency, instead of processing each date separately, five date blocks were supplied for the alignment and initial sparse point cloud formation steps establishing a common geometry between dates [Feurer and Vinatier, 2018]. Details of this processing are given in Section S2.

Hyperspectral preprocessing is described in detail by Laybros et al. [2019] and summarised here. Two sensors mounted to an aircraft side-by-side were used to cover the full 416–2500 nm wavelength range (see Table 1, Fig. S3). To merge the data from the two hyperspectral sensors without degrading the spatial resolution of the visible–near-infrared (VNIR) imagery, we resampled the short-wave infrared (SWIR) imagery to 1 m using nearest-neighbour interpolation. Images were orthorectified and georeferenced at 1 m spatial resolution with the PARGE software using a canopy Digital Surface Model (DSM) derived from a lidar acquisition contemporaneous with the 2016 hyperspectral campaign, so that the surface model used for orthorectification matched the canopy state at the time of the flight (Section S2). Bands in the SWIR with a low signal to noise ratio due to water absorption peaks were removed leaving 378 of the 448 total bands, and per pixel illumination was calculated using the shadow detection method of Schläpfer et al. [2018]. Spectra were extracted from the overlapping flight lines rather than from a mosaic, retaining multiple views of individual crowns, which improves classification performance [Laybros et al., 2019]. Brightness normalisation was then applied to each pixel spectrum, dividing each band’s reflectance by the sum across all bands [Dalponte et al., 2014]. Because classifiers sensitive to feature scale are otherwise dominated by bands with higher absolute variability, we also applied ‘standard’ scaling, removing the mean and scaling to unit variance (see Fig. 6). Additional details, including co-registration, are given in Section S2.

### 2.4 Temporal compatibility of data sources

The remote sensing datasets span different acquisition dates (Table 1): UAV RGB imagery from 2020–2021, hyperspectral imagery from September 2016, and the lidar CHM used throughout the analysis from November 2019. Each crown polygon carries validity dates recording when mortality, major branch fall, or significant crown change was observed, so crowns can be filtered to the epoch of each sensor; of the 3,500 crowns in the database, 3,256 had valid species assignments and sufficient temporal overlap with the 2016 acquisition for the classification task. Segmentation used only crowns valid during the 2020–2021 RGB campaign and classification only crowns valid during the 2016 hyperspectral acquisition, so each stage is trained and assessed against imagery matching the canopy state it represents. Section S2.4 sets out the distinct roles of the two lidar acquisitions and the evidence that crown shape is stable over the intervening intervals.

### 2.5 Field-derived tree crown database

To train and validate our models we generated a set of hand delineated, ‘ground truth’ crowns with species labels, built and curated between 2015 and 2023 and validated over eight field missions. Crowns were delineated in QGIS on the lidar CHM, which provides the greatest stability and spatial precision for outlines, with RGB, multispectral and hyperspectral layers overlaid to exploit as much spectral, textural and shape information as possible (see Table 1). Where crowns fell within inventory plots, an initial match to an individual was assigned from trunk location and size. Two provisional confidence scores were recorded: a ‘crown integrity’ score, describing the certainty that the outline defines the complete crown of a single individual rather than a partial crown or several crowns, and a ‘trunk match’ score for the confidence of assignment to an inventory individual. Changes through mortality or branch fall were date-stamped so crowns could be filtered to match the remote sensing source they are paired with. Subsequent fieldwork refined the delineations: comparing *in situ* observations with the imagery, we either matched crowns to trees in the site inventory [Gourlet-Fleury et al., 2004] or, for crowns outside the known plots, engaged botanists to assign species. Outlines and confidence scores were updated accordingly and liana infestations noted. Unlike previous studies [Laybros et al., 2019], infested crowns were retained and flagged throughout. Additional details are in Section S3.

### 2.6 Consensus mapping of tree crown locations through repeated surveys

The field-delineated tree crowns were partitioned into a training and testing set based on their geographic location (see Fig. 1). Regional partitioning ensured clear spatial separation between training and testing datasets, thus negating potential inflation of reported accuracy induced by spatial autocorrelative effects (see Kattenborn et al. 2022).

We used *detectree2*, a tool based on the Mask R-CNN deep learning architecture for automated tree crown delineation [Ball et al., 2023], which has been shown to outperform another leading CNN method for tree crown detection [Gan et al., 2023]. *Detectree2* was trained on the manual crown delineations and corresponding RGB images from the training dataset (see Fig. 1). Given that researchers seeking to map trees in their landscape of interest may or may not have some ground truth crowns and RGB surveys to train a model on, we opted to test models trained under different regimes:

1. The first was a ‘base’ model. This was not trained on the UAV RGB imagery and was just exposed to the plane mounted data and crowns from the range of sites described in Ball et al. [2023]. This meant it had been exposed to the Paracou forest but with a different sensor, different resolution imagery and four years separation. This pre-existing model is openly available for anyone to use^1^.
2. The ‘1 date’ model took the ‘base’ model and further trained the model on just the first date of the UAV RGB imagery and manual crowns.
3. The ‘5 date’ model took the ‘base’ model and further trained the model on the first five dates of the UAV RGB imagery and manual crowns.

Comparing the performance of these models provides an idea of what level of accuracy would be expected for researchers aiming to map their landscape with differing amounts of training data.

To test our consensus mapping approach, the trained models detected and delineated crowns across the entire UAV RGB region for all 10 acquisitions between 23-Oct-2020 and 06-Apr-2021. Images were tiled, predicted upon and recombined to give a set of crown polygons per date (see Ball et al. 2023 for the detection method), each carrying a confidence score (0–1). Where predictions overlapped (Intersection-over-Union, IoU ≥ 0.2), the most confident was retained. Per-date predictions were then merged into *consensus* delineations, from two dates up to the full ten, to determine the marginal benefit of each additional survey.

The consensus-fusion procedure operated in three stages (full mathematical detail and parameter settings in Section S4). First, all per-date predictions were pooled and pairwise IoU computed, with polygons at IoU ≥ 0.75 grouped as candidate observations of the same physical crown. Second, within each group, boundaries were resampled to 300 vertices, aligned by a common starting point (minimum *x* + *y*), and averaged using confidence scores as weights, with the fused polygon scaled to the mean area of its constituents; each consensus crown carried the summed confidence *C*_sum_ and the number of dates on which it was detected. Third, consensus polygons were sorted by descending *C*_sum_ and greedily placed into the landscape, accepted only where IoU with all already-placed polygons was < 0.2, yielding a non-overlapping, temporally stable crown map. All thresholds (cross-date match IoU ≥ 0.75; placement conflict IoU ≥ 0.2; quality filter *C*_sum_ > 0.1 × *n*) were tuned on training crowns and fixed for evaluation.

To evaluate the performance of the segmentation algorithm, we measured the overlap between predictions and reference crowns. An IoU of an overlapping pair of more than 0.5 was considered a match. This is a standard threshold used in the comparison of tree crown segmentation algorithms [Aubry-Kientz et al., 2021] that allows for small discrepancies in alignment and outline. These true positives, as well as the unmatched predictions (false positives) and unmatched manual crowns (false negatives), were used to calculate the precision, recall and F1-score of the predictions. To determine whether combining tree crown segmentation predictions across dates improved the segmentation accuracy through consensus building, we assessed the F1-score of each combination of dates, from each single date prediction to the combination of all ten dates. By taking all possible date combinations, we estimated a mean and standard deviation for the F1-score for each level of multi-date combination (single date through to ten dates).

### 2.7 Representative tree species classification

Of the 3,500 manual crowns in the database, 3,256 were labelled to species level (239 species) with sufficient confidence for classification. Of these, 169 species had at least two crowns, allowing at least one crown for training and one for testing. The 70 singleton species were excluded from model training and testing, but their proportional representation was retained in the weighted F1 calculation (using frequencies from the original, unfiltered dataset) to avoid inflating landscape-level accuracy estimates. Training and prediction used individual pixels (each labelled with species and containing reflectance values for 378 bands) rather than whole crowns, to accommodate species with few crowns and to improve spatial transferability across varying atmospheric and illumination conditions. Pixels with modelled illumination fraction below 60% (estimated from the lidar-derived DSM and solar geometry; Schläpfer et al. 2018) were discarded to exclude heavily shadowed areas. The successive filters that reduce the full crown database to the samples used for classification and for the repeated cross-validation are summarised in Table 2.

**Table 2:** Data flow from the full field-derived crown database to the samples used in the classification analyses. Each row applies the stated selection criterion to the preceding row; the final row reflects the crowns retained after per-pixel illumination and crown-footprint filtering that are used in the 20×5-fold repeated cross-validation.

| Stage | Crowns | Species | Selection criterion |
| --- | --- | --- | --- |
| Delineated crown database | 3,500 | – | Manually delineated, field-curated (2015–2023) |
| Species-labelled pool | 3,256 | 239 | Labelled to species with sufficient confidence |
| Classification set | 3,186 | 169 | Species with $\geq 2$ crowns (70 singleton species removed) |
| Repeated-CV set | 2,762 | 169 | Retained after illumination ($< 60\%$) and crown-footprint filtering (317,097 pixels) |

Whereas crown delineation required regional partitioning between train and test sets, species identification was better served by species-stratified, crown-level partitioning. Given the diversity and mixing of the forest, the average distance between conspecific crowns was large enough not to require additional spatial constraints controlling for autocorrelation (cf. Wadoux et al. 2021, who argue that spatial CV can itself introduce bias when clusters lack a statistical basis; grouping by individual crown provides a natural, non-arbitrary unit that prevents within-crown autocorrelation leaking into accuracy estimates). A test set of a stratified random 20% of crowns per species was therefore withheld until the final evaluations, except where a species had four or fewer individuals, in which case a single test crown was drawn at random. Species with a single recorded individual were excluded.

To test which machine learning classifier yielded the best overall accuracy, models were trained on hyperspectral data extracted from the delineated crowns, spanning 378 bands, 169 species, 99 genera and 41 families. In line with common practice we evaluated Multi-Layer Perceptrons (MLPs), Linear Discriminant Analysis (LDA), Random Forest (RF), Support Vector Machine (SVM), k-Nearest Neighbours and Logistic Regression. Further approaches – dimensionality reduction (PCA, UMAP), pipeline combinations (e.g. PCA+SVM), gradient boosting (HistGradientBoosting, LightGBM, XGBoost), PLS Discriminant Analysis, elastic-net logistic regression, and covariance-regularised variants (LDA with Ledoit-Wolf shrinkage, Quadratic Discriminant Analysis (QDA)) – were explored but not taken forward as they failed to improve accuracy (Section S5.2). We deliberately retained all 378 bands without feature selection, because LDA performs its own implicit dimensionality reduction by projecting into discriminant space; because retaining all bands allowed the ablation analysis (Section 2.8) to quantify band importance directly; and because we sought a broadly reproducible approach, avoiding feature engineering that may not transfer to other sites or sensors. Training on individual pixels left the data heavily imbalanced: crowns contained 115 pixels on average (317,097 pixels across 2,762 crowns), ranging from 55,448 pixels for the most abundant species to 223 for the rarest. Pixel resampling and inverse-frequency class weighting were tested but degraded overall accuracy (Section S5.2), because down-weighting abundant species sacrifices signal that supports landscape-representative mapping; we therefore retained the natural prevalence distribution. All metrics were computed at crown level via pixel-wise majority vote, so each test crown contributes once and large crowns cannot dominate the evaluation. Pixel-level training nonetheless risks leakage under naïve cross-validation, since adjacent pixels within a crown share local effects; we therefore used a stratified-group 5-fold cross-validation in which all pixels from a given crown fell exclusively in either the training or validation partition. By design, the resulting maps are most accurate for the common canopy species that dominate landscape area, a point we return to in the Discussion.

After cross-validation, models were retrained on the full training set and used to predict species across the landscape, with crown-level classifications determined by majority vote of constituent pixels. Final performance on the held-out test set was evaluated using *weighted F1* (F1 averaged across species and weighted by each species’ proportional abundance in the reference dataset, giving a landscape-representative estimate), *macro-average F1* (the unweighted mean across species, treating each equally regardless of abundance), *crown-level accuracy* (the fraction of test crowns assigned the correct species), and the analogous *weighted precision* and *weighted recall*. We use these terms consistently throughout.

**Robustness assessment via repeated cross-validation.** To obtain per-species estimates that are not contingent on a single train–test split, we additionally ran a 20-times-repeated stratified-group 5-fold cross-validation over 2,762 crowns (169 species; 317,097 pixels) retained after illumination and footprint filtering, again grouping by crown to prevent leakage. LDA was trained on each fold’s training pixels and evaluated at crown level via majority vote, yielding per-species F1 scores for every fold and 100 estimates of the number of species exceeding F1 ≥ 0.7, from which we derived an empirical 95% confidence interval on that count. As a complementary single-split characterisation of uncertainty, we also computed Bayesian posterior median F1 scores with 95% credible intervals for each species on the held-out test set, using a Dirichlet posterior over the confusion-matrix entries [Tötsch and Hoffmann, 2021].

To assess whether including rarer species in training is detrimental to overall accuracy, we reran the best-performing model on a series of artificially restricted datasets, iteratively removing individuals of the least well-represented species until only two species remained (Section S5.3). Two scenarios were examined: (i) the test pool was kept complete (169 species) while the training pool was progressively restricted; and (ii) the test pool was reduced in parallel with the training pool.

### 2.8 Evaluate which wavebands are most important for distinguishing species

To determine which wavebands were most important for distinguishing species, we extracted feature importance from the LDA classifier using variance-weighted squared discriminant scalings [Peterson and Mahajan, 1976]. For each discriminant axis *k*, LDA produces a vector of scalings *w_k_* defining the linear projection that maximises the ratio of between- to within-class variance. The importance of band *j* was the sum of its squared scaling across all *K* axes, weighted by each axis’s explained variance ratio, 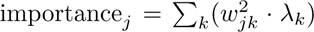, normalised to sum to one; this is sign-invariant and gives greatest weight to the axes contributing most to class separation (Section S6.1).

Because scalings measure each band’s *unique* contribution after controlling for all others, they can be sensitive to multicollinearity among the 378 channels. To assess the stability of the rankings, we extracted scalings from the LDA models trained in each of the 100 cross-validation folds and computed per-band importance across all folds. As a complementary perspective we also computed variance-weighted squared structure coefficients [Courville and Thompson, 2001] – the correlation between each band and the discriminant scores – which capture each band’s *total* association with species discrimination and are robust to multicollinearity (Section S6.1).

To test how these importances transferred to prediction accuracy we performed ablation experiments, progressively removing bands in increments of ten by setting their standardised values to zero before training. Four removal orders were used: most to least important; least to most important; random shuffling of 15 k-means clusters of band importance, which retains band order within clusters and so preserves a realistic data structure given that importance tends to cluster in specific regions; and fully random shuffling, which leaves all spectral regions available for longer than the other procedures. See Section S6.2 for details.

## 3 Results

### 3.1 Tree crown segmentation using multiple surveys

Tree crown segmentation accuracy was improved by combining multiple dates of tree crown segmentation predictions and retaining crowns that had good confidence and agreement between dates (see Fig. 3). We compared the accuracy of segmentation from using a single time step and multiple time steps with an unseen test set of 169 test crowns across two spatially separate test zones. The accuracy of delineations increased as more dates were combined. The best performing model overall was the one trained on the broadest range of imagery (five individual dates’ worth). The accuracy of its delineation was boosted significantly by combining the delineations from different time steps (consensus mapping) from a mean F1-score of 0.68 for a single date prediction, to 0.78 when combining nine to ten time steps (see Fig. 3). In terms of total crown area, approximately 86% of the test region had well located and delineated crowns (see Fig. 4). Accuracy tended to increase with tree crown area.

**Figure 3:**
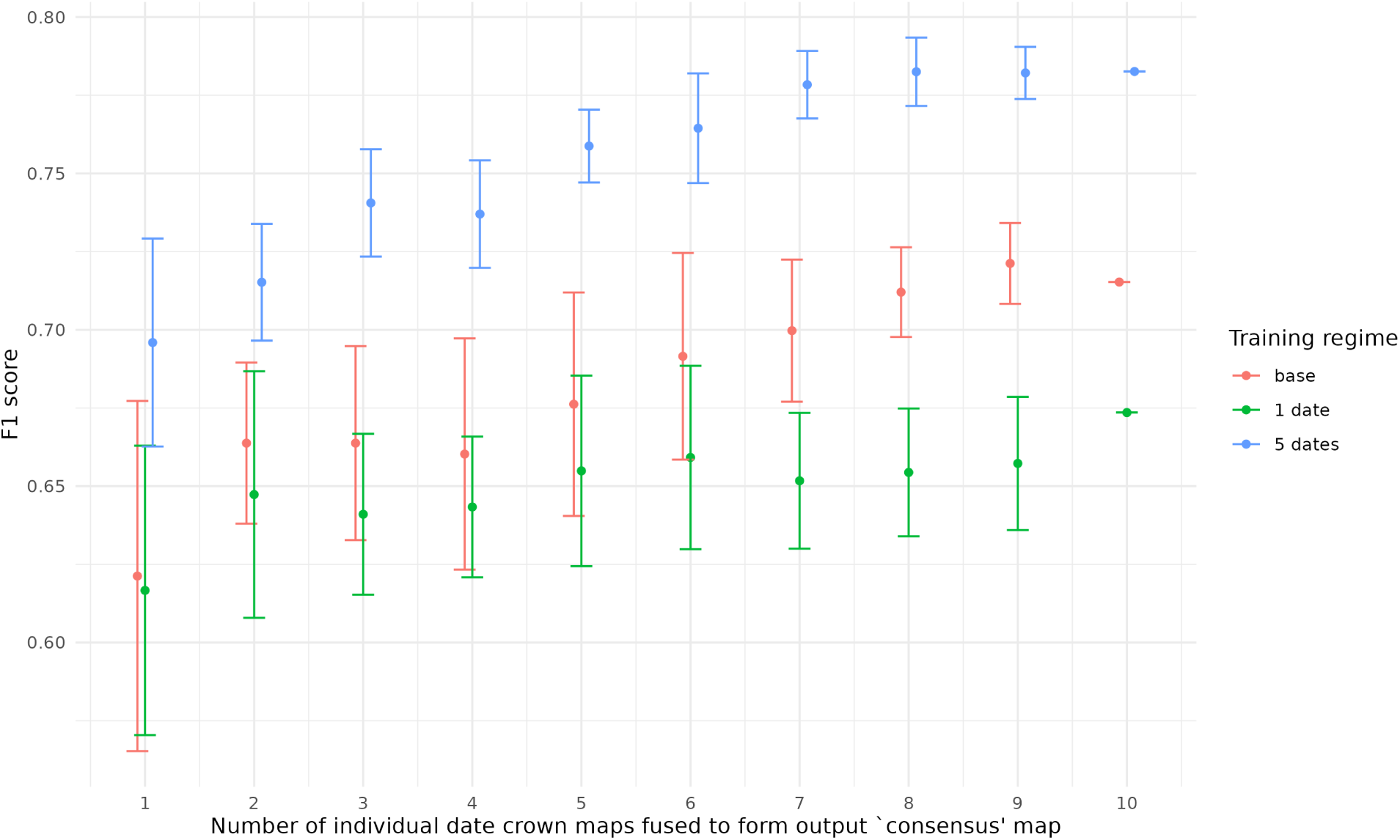
Crown segmentation F1-score as a function of the number of date-maps combined via consensus fusion, evaluated on 169 test crowns across two spatially separate test zones. Three models were tested: (1) the ‘base’ model, freely available online and trained on non-UAV imagery from multiple sites; (2) the ‘1 date’ model, the base model fine-tuned on a single UAV-RGB survey; (3) the ‘5 date’ model, the base model fine-tuned on five UAV-RGB surveys. Points show the mean F1-score across all possible date combinations at each combination level; error bars show ±1 standard deviation across these combinations (not statistical confidence intervals). At the 10-date level only a single combination exists.

**Figure 4:**
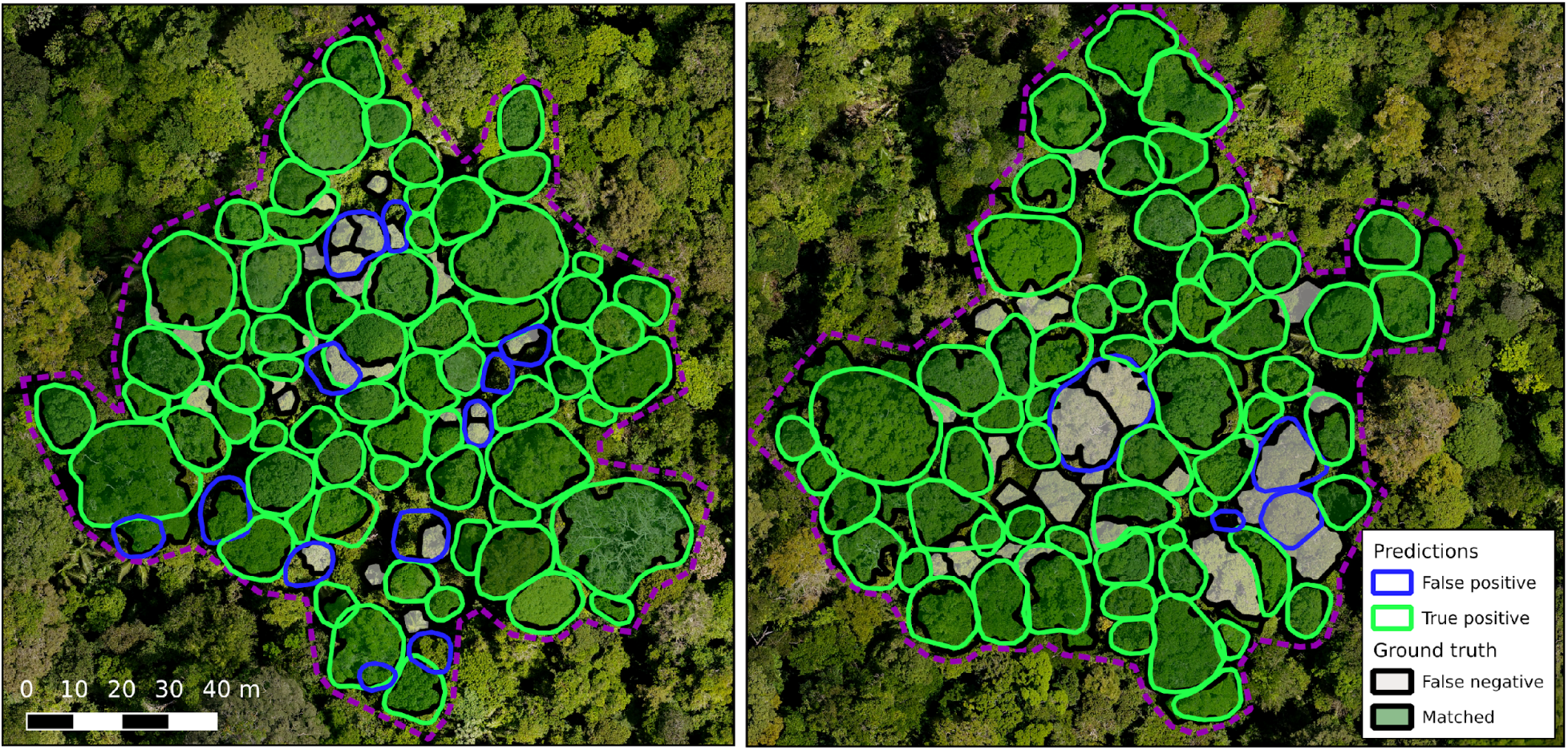
Predicted and reference crowns in the two unseen test zones (10-date consensus map, 5-date model). Reference crowns: black border, green fill if matched to a prediction (IoU ≥ 0.5), grey fill if unmatched. Predicted crowns: green border if matched to a reference crown, blue border if unmatched (false positive).

The ‘base’ model and the ‘1 date’ model performed comparably for single-date predictions and for combinations of fewer than seven dates, but beyond seven the ‘base’ model became substantially better, surpassing the single-date accuracy of the model trained on five dates (Fig. 3). The ‘1 date’ model gained less from combining dates than the other two, suggesting overfitting: trained on a range of non-UAV RGB imagery without focused UAV training [Ball et al., 2023], the base model achieved better temporal transferability than focused training on a single date.

### 3.2 Tree species classification using hyperspectral imagery

The field teams labelled 3,256 crowns to species level with sufficient confidence, comprising 239 species, 124 genera and 43 families. We removed 70 species encountered only once, as they could not be independently trained and tested, leaving 169 for analysis (Fig. 5). Crowns per species were highly skewed: median 6 (interquartile range 2–13), from 2 crowns for the rarest included species to 223 for the most abundant.

**Figure 5:**
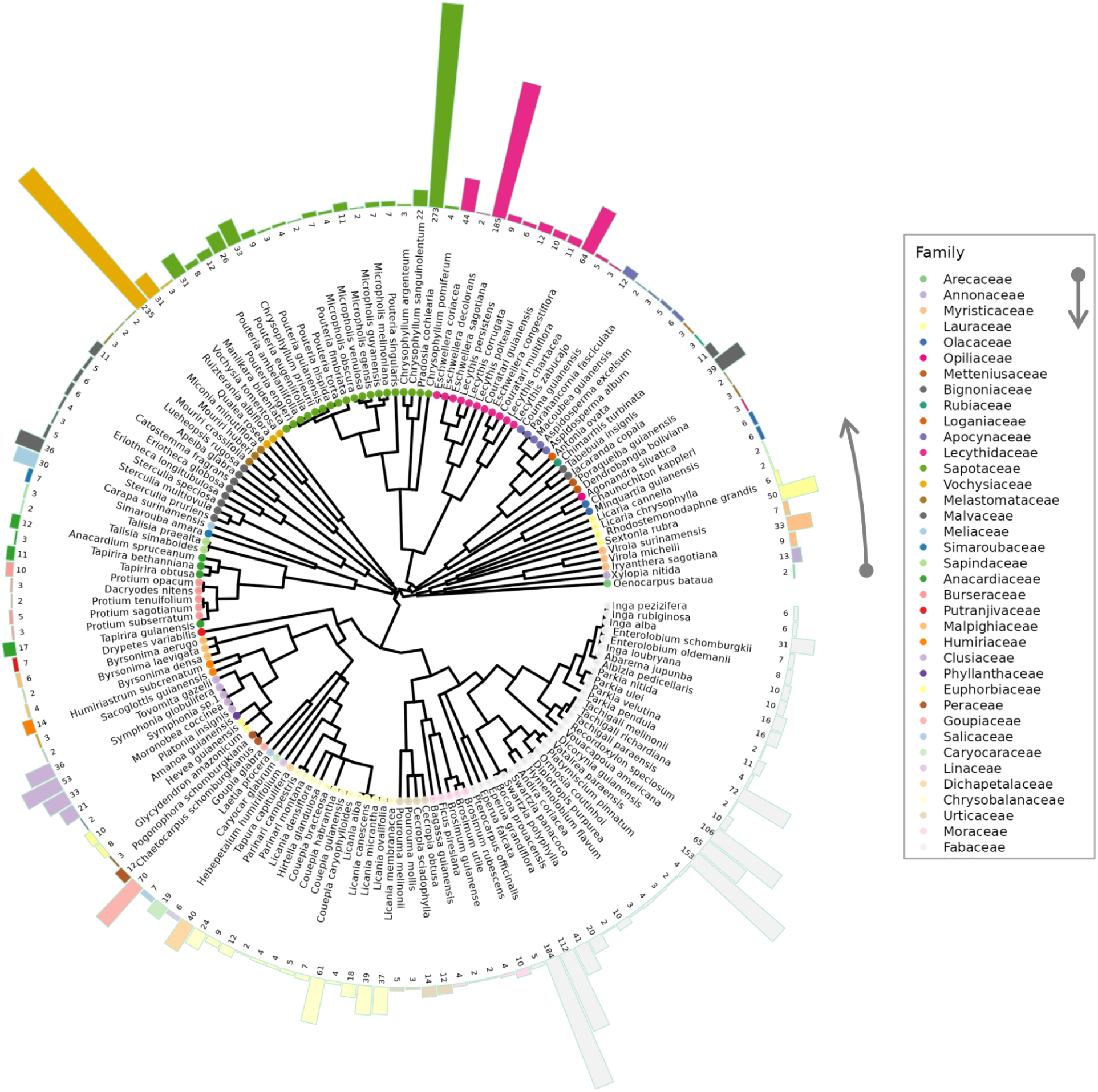
Abundance of 169 species in the field-validated crown dataset, mapped onto the phylogeny of Baraloto et al. [2012]. Bar length indicates the number of crowns per species; colours denote taxonomic family (legend ordered as families appear anti-clockwise from 3 o’clock). The 70 singleton species excluded from classification are not shown. An enlarged version with fully legible species labels is provided in the online supplementary materials.

The LDA classified test crowns best from their hyperspectral signal (weighted F1 = 0.75; Table 3; confusion patterns in Fig. S11). Logistic regression performed slightly worse overall but with a substantially lower macro-average F1, suggesting it struggled with less well-represented classes. The more flexible, and more expensive, MLP and SVM classifiers failed to match the LDA, indicating that its less flexible approach to separating classes transferred more robustly between crowns. The LDA was also far quicker, training in 20 seconds against more than 7 hours for the SVM, whose quadratic-to-cubic complexity in the number of training samples makes it costly on this many pixels^2^.

**Figure 6:**
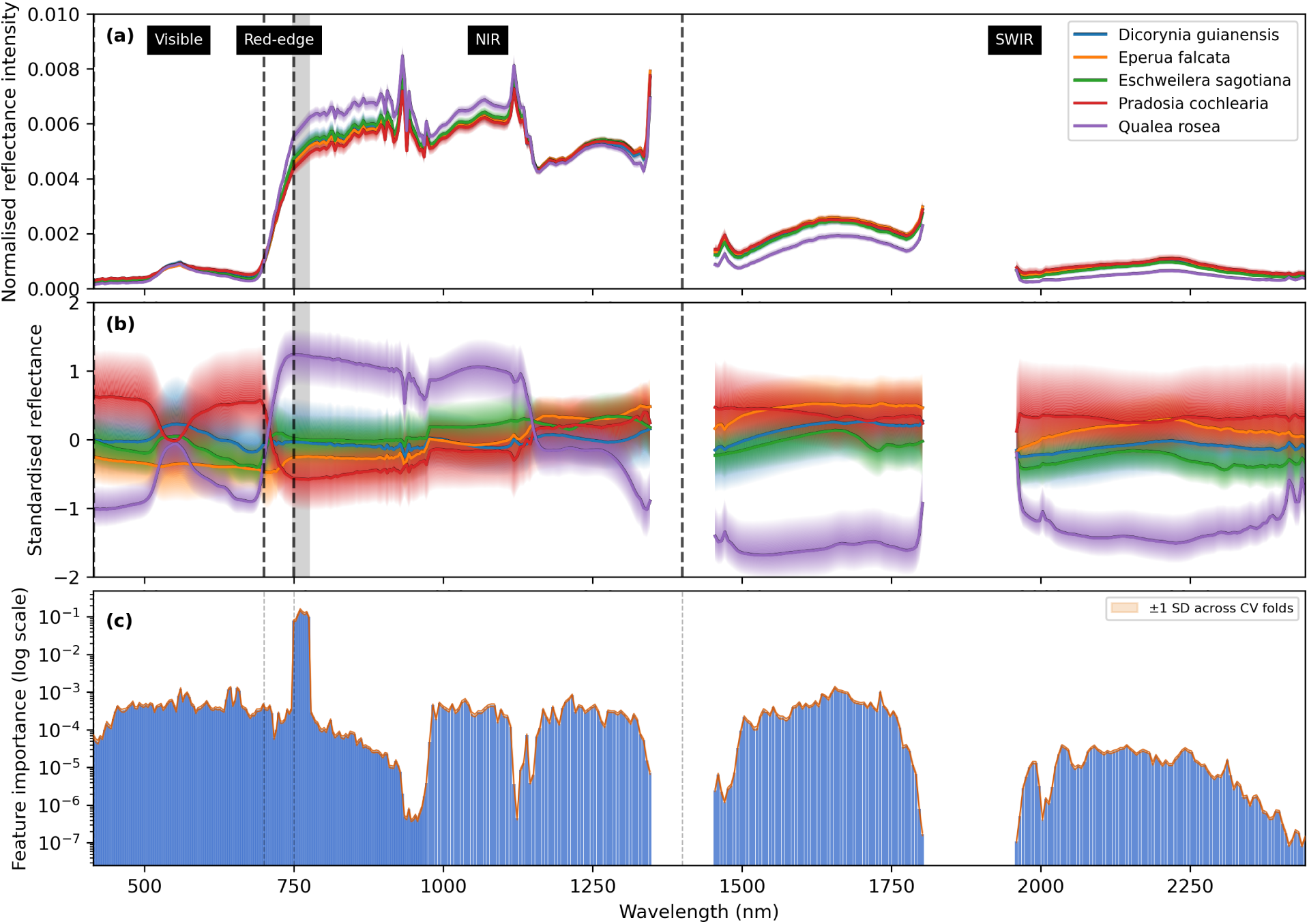
Spectral signatures of five common species and per-band importance for classification. Lines show the median reflectance per band across crown pixels; shading shows the interquartile range. (a) Normalised reflectance (each pixel spectrum divided by its total reflectance). (b) Standardised reflectance (per-band mean subtracted and divided by the per-band standard deviation across valid crown pixels), allowing comparison on a common scale. (c) LDA band importance (best model, trained on the standardised data): taller bars indicate greater contribution to interspecific separation. The grey vertical band highlights the far-red edge (748–775 nm), identified as most important for species classification. SWIR gaps correspond to bands removed because of atmospheric humidity effects.

**Table 3:** Classification performance on the held-out test set (169 species). Weighted metrics use species abundances from the full crown dataset (prior to singleton removal and train–test splitting) so that landscape-level accuracy is not inflated by the exclusion of rare species. Crown-level accuracy is the fraction of test crowns assigned the correct species via majority vote of constituent pixels. Train time is for the final tuned model on a 128-core 2×AMD EPYC 9534 with 1.5 TB RAM and NVIDIA A30 GPU.

| Classifier | Weighted precision | Weighted recall | Weighted F1 | Macro-average F1 | Crown-level accuracy | Train time |
| --- | --- | --- | --- | --- | --- | --- |
| **LDA** | **0.74** | **0.79** | **0.75** | **0.49** | **0.78** | **20s** |
| MLP | 0.65 | 0.74 | 0.68 | 0.37 | 0.72 | 2m26s |
| Logistic | 0.66 | 0.74 | 0.68 | 0.35 | 0.72 | 45m20s |
| SVM | 0.64 | 0.71 | 0.65 | 0.33 | 0.70 | 7h11m6s |
| QDA | 0.52 | 0.58 | 0.53 | 0.21 | 0.57 | 14s |
| Random Forest | 0.41 | 0.43 | 0.35 | 0.11 | 0.42 | 41s |
| kNN | 0.35 | 0.39 | 0.31 | 0.08 | 0.38 | 3s |

Using the best-performing classifier, 81% of the total crown area in the test set was correctly identified to species level. Combined with the crown area that was well located and delineated (86%), we conservatively estimate that 70% of the landscape’s crown area was mapped correctly. This assumes that accuracy on manually delineated crowns is representative of performance on automatically segmented ones; because classification operates on pixels within crown boundaries, the assumption is reasonable provided crowns are correctly located and delineated, though end-to-end validation of the fully automated pipeline would strengthen the claim. For reference, the twenty most abundant species make up < 60% of the reference dataset’s total crown area (Fig. 7).

**Figure 7:**
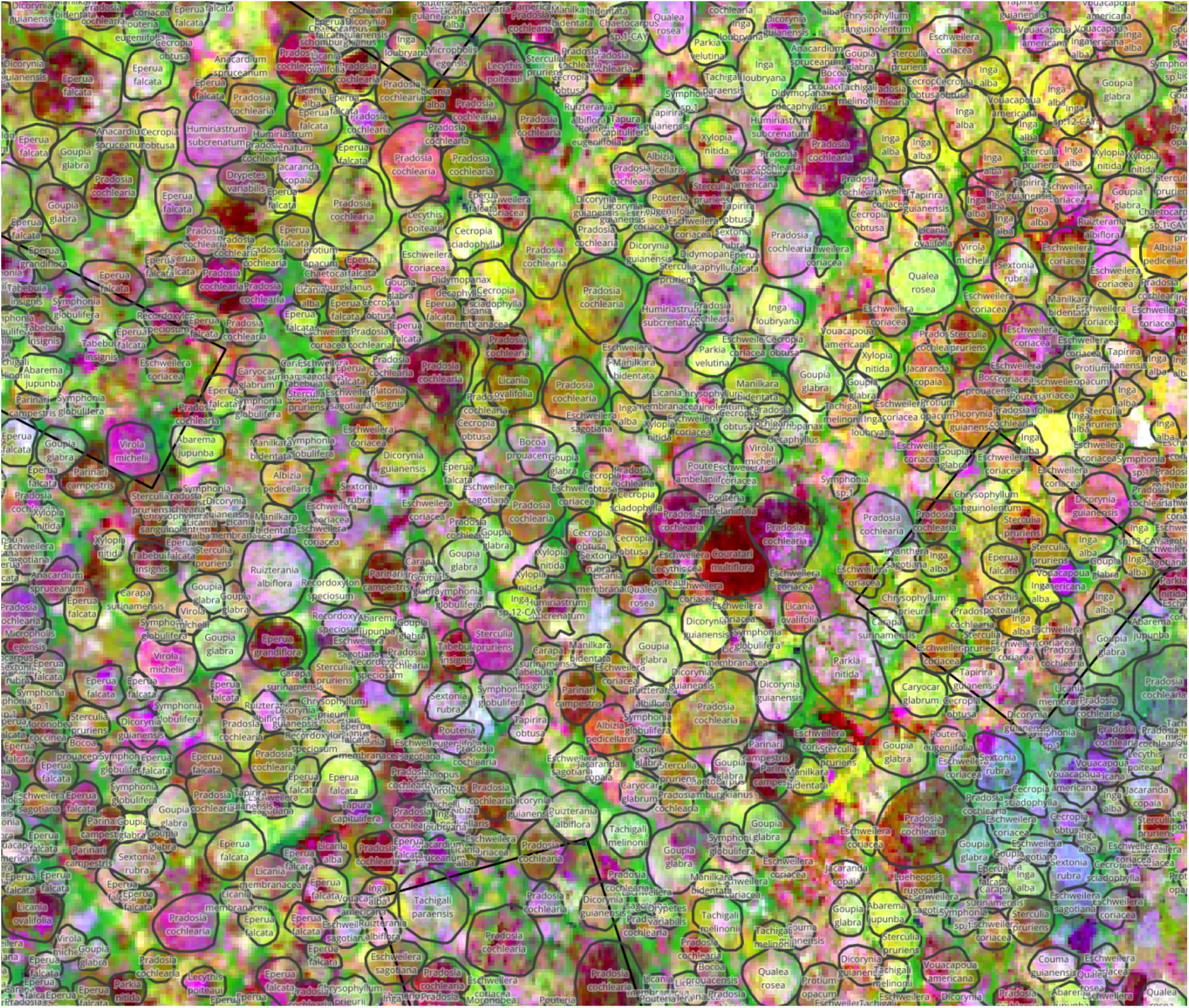
Final species-labelled crown map over a hyperspectral PCA backdrop. Background shows an RGB composite of three principal components (PC1–PC3) from the hyperspectral imagery for visualisation only (PCA outputs were not used in the analysis). Predicted crowns are outlined in black and labelled with species; black squares denote forest inventory plots. Inset panels show zoomed regions with legible species labels.

**How many species can be reliably classified?.** The aggregate weighted F1 of 0.75 masks a strongly bimodal distribution of per-species performance. To assess how many species can genuinely be classified at F1 ≥ 0.7, and how sensitive that count is to the particular train–test split, we ran a 20-times-repeated stratified-group 5-fold cross-validation over 2,762 crowns (Section 2.7). Across 100 folds, the number achieving crown-level F1 ≥ 0.7 in a given fold averaged 50 (95% CI: 41–63; Fig. 8 inset), with a weighted F1 of 0.72 (SD 0.02) and macro-average F1 of 0.48 (SD 0.03) confirming stable aggregate performance. Averaged across folds, 38 species maintained a mean F1 ≥ 0.7 and 15 exceeded the threshold in at least 80% of folds – a core group we consider reliably classifiable. The single held-out test set corroborated this: 67 of the 169 species achieved point-estimate F1 ≥ 0.7, though Bayesian credible intervals showed only 25 had lower bounds above 0.5, the remainder having too few test crowns for precise estimation. That figure of 67 sits at the upper bound of the CV-derived 95% CI, so it is plausible but likely optimistic relative to expected performance on new data.

**Figure 8:**
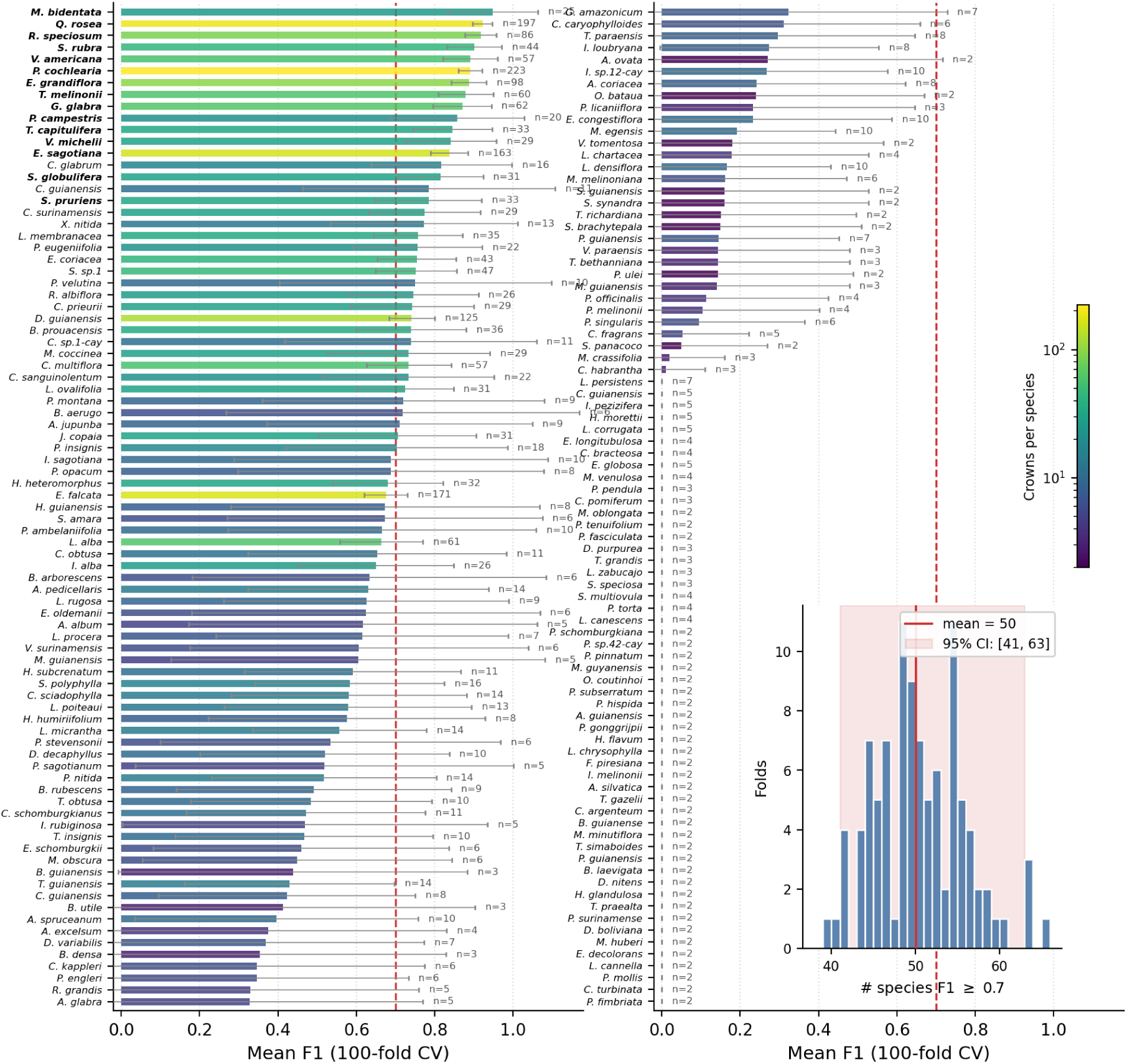
Per-species classification performance from repeated cross-validation (LDA, 169 species). Each point shows the mean F1 across 100 CV folds (20 repeats x 5-fold stratified-group, grouped by crown); vertical bars indicate ± 1 SD. Points are coloured by the number of crowns available per species (log scale). The dashed line marks F1 = 0.7. Of the 169 species, 38 achieved mean F1 ≥ 0.7 and 15 exceeded this threshold reliably (in ≥ 80% of folds; species names in bold). Inset: distribution of the number of species achieving F1 ≥ 0.7 across folds; the solid line marks the mean (50), shaded band the 95% CI (41–63).

These numbers are clearer expressed as canopy area. The 38 species with mean CV F1 ≥ 0.7 account for approximately 70% of total mapped canopy area (206,000 of 296,000 m^2^), and the 15 reliably classified species alone cover 48%. Because common species dominate canopy area in hyperdiverse forests, even a modest number of well-classified species translates to substantial landscape coverage, while the many hard-to-classify rare species contribute relatively little area. Per-species F1 was positively associated with the number of available training crowns (Fig. 8); the formal analysis of this and other determinants is presented in our companion paper [Ball et al., 2026].

**Inclusion of rare species.** To test whether rare species in the training pool degrade classification of better-represented species, we progressively removed the least-represented species from training while retaining the full 169-species test set (scenario i; Section S5.3). Contrary to expectation, weighted F1 decreased as species were removed (Fig. S4), indicating that including rare species imposes no penalty on landscape-level accuracy and may even provide ancillary discriminative information. Under scenario (ii), where the test pool shrank in parallel, accuracy rose as expected given the reduced task. These results support retaining all available species in training.

### 3.3 Which spectral regions are most important for distinguishing species?

Eight bands between 748 and 775 nm dominated in terms of relative feature importance for separating species (Fig. 6c). We refer to this narrow window on the near-infrared shoulder, just beyond the conventional red-edge inflection point (700–750 nm), as the far-red edge (FRE). Using variance-weighted squared scalings, the FRE accounted for 91.5% ± 0.3% of total importance (mean ± SD across 100 CV folds), and importance rankings were extremely stable across folds (mean pairwise Spearman rho = 0.997). The same eight FRE bands appeared in the top ten in every fold (95.4% overlap with the overall ranking). The next most important regions were 640–660 nm (red), 560–575 nm (green), 1630–1680 nm (SWIR), and 1000–1100 nm (NIR).

The degree of FRE concentration depends on how importance is measured: it reflects the concentration of *unique* (non-redundant) discriminative information in the scalings-based metric. When measured via structure coefficients – which capture the *total* correlation between each band and species discrimination, including information shared with other bands – importance was distributed much more evenly across the spectrum, with the FRE contributing only 2.5% (Fig. S5). These two perspectives are complementary: scalings identify where the classifier concentrates its discriminative power, while structure coefficients reveal that species-relevant spectral information exists broadly across the 416–2500 nm range (see Section S6.1 for full comparison).

Ablation tests validated the scalings-based ranking. Removing the ten most important bands caused no immediate, dramatic drop in performance, as the remaining bands partially compensated (Fig. S6), but progressive ablation from most to least important degraded performance substantially faster than reverse or randomised orders. Fully randomised removal preserved performance longest, leaving all spectral regions partially available until late in the process because adjacent bands share information. These results confirm that the FRE concentrates the most non-redundant discriminative information, while classification also draws on complementary cues across the full spectral range (Section S6.2).

## 4 Discussion

### 4.1 Overview

Our results indicate that species-level mapping of tropical forest canopies at landscape scale is achievable at substantially greater scope than previously reported, but with important caveats. While earlier airborne hyperspectral studies in the wet tropics typically resolved 20 or fewer common canopy species [Féret and Asner, 2013, Laybros et al., 2019, Garzon-Lopez and Lasso, 2020], repeated cross-validation showed that on average 50 species achieve F1 ≥ 0.7 in any given fold (95% CI: 41–63), with 15 doing so reliably across folds (Section 3.2); those 15 alone account for roughly 48% of mapped canopy area in a forest where more than one hundred tree species per hectare are commonly found [Lee et al., 2002, Bohlman, 2015, Duque et al., 2017]. However, even the upper-bound estimate of ∼67 species is less than 10% of the more than 800 species (including sub-canopy species) recorded across the Paracou plots, and many rare species remained effectively unclassifiable. This advance rests on four interacting factors specific to our study: a dense, well-curated reference dataset of geo-located, confidently identified crowns built over a decade of fieldwork; precise field-to-image co-registration; high-resolution hyperspectral imagery; and improved crown segmentation via a CNN with temporal consensus mapping. Whether comparable results can be obtained at other sites, with different sensors, or across larger extents remains to be tested.

### 4.2 Advances in Crown Segmentation

Temporal consensus mapping improved crown segmentation even when using a model trained elsewhere, showing that ensembling predictions across dates can substitute for limited local training data. We used *detectree2* [Ball et al., 2023], a Mask R-CNN–based model [He et al., 2017] that integrates spectral, textural, and geometric cues; trained on carefully validated field-mapped crowns it achieved state-of-the-art performance [Gan et al., 2023], even surpassing human annotators in complex canopies (Section S1). The key innovation was temporal fusion. Notably, the base model – trained on diverse imagery including earlier plane-mounted acquisitions over Paracou but no data from this UAV campaign – matched or exceeded models fine-tuned on a single local UAV date once predictions were aggregated (Fig. 3). We attribute this to overfitting: fine-tuning on one date specialises the model to that acquisition’s illumination and canopy state, whereas the base model’s broader training distribution provides implicit regularisation that yields more temporally robust features, an advantage amplified by multi-date consensus. This is, to our knowledge, the first demonstration in tropical forests that multi-date aggregation improves segmentation accuracy, offering a scalable route to monitoring forest structure, growth, mortality, and phenology across large landscapes.

### 4.3 Species classification accuracy: achievements and limitations

The weighted F1 of 0.75 across 169 species represents a substantial advance over previous tropical hyperspectral studies. However, the macro-average F1 on the same held-out test set is only 0.49, revealing that this performance is heavily driven by the most common species. The spread of per-species performance on that test set was wide: 67 of the 169 species reached F1 ≥ 0.7, 35 fell between zero and that threshold, and the remaining 67 – rare species with few training crowns – were never classified correctly. This pattern is expected where the abundance distribution follows a long tail: a minority of dominant species account for most canopy area, while most species are represented by a handful of individuals [ter Steege et al., 2013]. The weighted metric therefore provides the most meaningful measure of practical mapping accuracy – what proportion of the canopy is correctly labelled – while the macro-average highlights the challenge that remains for rare species. Repeated cross-validation (Fig. 8) reinforces this picture, and the species that are reliably classified are overwhelmingly the most abundant. Per-species performance is therefore better characterised as a continuum shaped by data availability than as a binary split between classifiable and unclassifiable taxa – a pattern explored in detail in our companion paper [Ball et al., 2026].

At this site, the relatively simple LDA consistently outperformed more flexible models such as SVMs and MLPs, echoing Féret and Asner [2013]. We interpret this as reflecting the high collinearity among the 378 spectral bands: LDA projects into at most *K −* 1 discriminant dimensions (here 168), effectively performing dimensionality reduction while estimating a single pooled covariance matrix shared across all classes. This implicit regularisation suits the high-dimensional, highly correlated structure of contiguous hyperspectral data, where the effective degrees of freedom are far fewer than the number of bands. Higher-capacity models, by contrast, can exploit fine-grained within-crown variation and tended to overfit to crown-specific artefacts, reducing transferability across individuals. The 20×5-fold repeated cross-validation, which explores 100 train–test configurations, confirmed that LDA’s advantage was not an artefact of a particular split. This result is nonetheless specific to this dataset’s spectral structure and should not be generalised to other sites, sensors, or tasks where non-linear decision boundaries may be more appropriate; it reinforces the need to constrain complex models – through regularisation, representation learning, or multi-instance methods – so they capture robust crown-level signals rather than artefacts.

Performance was driven primarily by the quality of the input data and preprocessing. The reference dataset of over 3,500 geo-located crowns was curated and field-validated over multiple years, and the labour-intensive image–field alignment yielded a rare, high-confidence resource. Such spatially explicit field datasets are essential for species-level ecological inference from remote sensing [Chave et al., 2019, Davies et al., 2021], yet remain scarce at this taxonomic and spatial resolution [Laliberté et al., 2020]. High-quality imagery further underpinned performance: acquisitions were near-nadir and precisely co-registered to crowns, enabling fine-scale analyses of trait-linked separability (Fig. 6). Our inclusive training approach, retaining rare taxa [ter Steege et al., 2013, Jackson and Adam, 2021], did not reduce landscape-level accuracy, suggesting training pools should remain as broad as possible (Fig. S4). Nevertheless, accurate identification of rare species remains difficult and will require targeted strategies such as few-shot learning and data augmentation.

### 4.4 Critical spectral regions for species classification

The far-red edge (FRE; 748–775 nm) emerged as the most important spectral region for species discrimination (Fig. 6), consistent with recent tropical findings [Badourdine et al., 2023]. While prior work broadly emphasised red-edge and NIR reflectance [e.g., Clark et al., 2005, Fassnacht et al., 2016, Hennessy et al., 2020], our analysis pinpoints a specific zone just beyond the conventional red-edge inflection of 700–750 nm [Guo et al., 2023]. This concentration of unique discriminative information was highly robust, with importance rankings nearly identical across 100 cross-validation folds (mean pairwise Spearman rho = 0.997), confirming it is not an artefact of a particular split or of multicollinearity among adjacent channels.

However, the apparent dominance of the FRE depends on how importance is defined. Scalings measure each band’s *unique* contribution after accounting for all other bands, whereas structure coefficients capture each band’s *total* correlation with the discriminant axes, including information shared with correlated bands [Courville and Thompson, 2001]; under the latter, importance was distributed far more evenly and the FRE contributed only 2.5% (Fig. S5). Together the two measures are coherent: species-relevant information exists broadly across the 416–2500 nm range, but the FRE uniquely concentrates its non-redundant component.

The FRE spans the transition between dominant chlorophyll absorption and internal leaf scattering in the NIR [Horler et al., 1983, Gitelson et al., 2003]. Reflectance here is shaped by pigment concentrations and by species-specific microstructural attributes – mesophyll architecture, refractive index gradients, and intercellular airspace – that control scattering efficiency, with leaf water content further modulating the balance through cellular turgor and optical path length [Boochs et al., 1990, Ustin et al., 2009]. That adjacent wavelengths score low on scalings but high on structure coefficients suggests they carry largely redundant information the FRE already captures. The FRE thus acts as a spectral “confluence” where biochemical, structural, and physiological axes intersect to amplify interspecific contrast, helping explain why species with broadly similar chlorophyll concentrations nevertheless diverge in this narrow range.

Secondary regions also contributed: 640–660 nm (red) likely reflects variation in chlorophyll b, 560–575 nm (green) may capture differences in leaf structure and carotenoid/anthocyanin content, and 1630–1680 nm (SWIR) corresponds to water absorption features and variation in cellulose/lignin composition. These signals, weaker than the FRE in the scalings metric, are proportionally more prominent under structure coefficients (Fig. S5), underscoring that species-relevant information is not confined to one region. Ablation showed that more than half of the 378 bands can be removed in randomised order before performance degrades substantially (Fig. S6), confirming redundancy among adjacent bands and suggesting that a reduced set – centred on the FRE with secondary coverage in the red, green, and SWIR – could retain most classification power. A comparison of model-agnostic feature selection strategies (Section S6.3; Fig. S7) confirmed this: MI-based selection reached F1 ≥ 0.70 with 200 bands and matched the full-band result by 300, while PCA-based reduction and random selection converged similarly or more slowly and never exceeded the full-band baseline. Across classifiers (Section S6.4; Fig. S8), LDA’s advantage over Random Forest and SVM *increased* with dimensionality – the opposite of overfitting – and 15 standard vegetation indices were insufficient for all classifiers (F1 ≤ 0.29). Prior feature selection is therefore unnecessary, and LDA’s superiority reflects its built-in regularisation rather than any artefact of high dimensionality. These rankings are, however, aggregate measures across all 169 species and may not reflect the optimal bands for any individual species or group.

### 4.5 Limitations and Future Directions

**Biological variability and timing.** Within-species spectral variability – arising from leaf ontogeny, physiological stress, liana infestation, crown position and microenvironment, and phenological stage – can obscure consistent species-level signals [Ollinger, 2011, Chavana-Bryant et al., 2019, Hesketh and Sánchez-Azofeifa, 2012], and even at a single time point individuals of the same species may spectrally diverge [Reich, 1995, Theiler et al., 2019, Badourdine et al., 2025]. Fully modelling the joint biochemical, structural and phenological controls with radiative-transfer frameworks remains intractable in hyperdiverse tropical canopies, because trait covariation and crown architecture are difficult to parameterise at scale [Viskari et al., 2019, André et al., 2021, Qi et al., 2022], which highlights the need for empirical, multi-temporal datasets and representation learning approaches that can absorb such variability. We retained liana-infested crowns rather than excluding them, since infestation is a common feature of tropical canopies that any landscape-scale map must contend with; our companion analysis [Ball et al., 2026] shows it lowers the odds of correct identification by approximately 38%, so excluding infested crowns would raise the headline figures at the cost of ecological realism.

Phenological and physiological asynchrony among individuals can shift the red-edge position – and thus the FRE – through changes in pigments and stress state [Curran, 1989, Gitelson et al., 2003, Clark and Roberts, 2012], reducing separability, especially when imagery is acquired during periods of low leaf area or poor synchrony [Hesketh and Sánchez-Azofeifa, 2012]. Conversely, well-defined cycles constrain within-species variance by aligning leaf developmental stages, and distinct events such as synchronous leaf flush or flowering can accentuate between-species contrasts [Takahashi Miyoshi et al., 2020]. Our dry-season acquisition coincided with substantial leaf flushing; future surveys could be timed to phenophases that maximise within-species alignment, improving spectral comparability and classification reliability.

**Multimodal classification.** Our approach focused on spectral separation, but additional data streams can aid species with overlapping spectra. Structural lidar metrics [Zhong et al., 2022, Quan et al., 2023], high-resolution texture [Williams et al., 2022], and multi-season imagery [Takahashi Miyoshi et al., 2020, Modzelewska et al., 2021] provide complementary information on crown architecture, leaf angle distribution and phenological cycling. Acquiring these at matched spatial and temporal fidelity across large or remote sites is, however, logistically demanding, and fusion increases alignment and model-complexity burdens – practical constraints for operational tropical monitoring.

**Addressing transferability.** The goal is classifiers that are robust across time and space, yet performance can degrade with modest shifts in acquisition conditions or community composition [Laybros et al., 2019]. Some errors – image/label misalignment, occasional annotation mistakes – are tractable; others – atmospheric effects, geometry, illumination – require stronger radiometric and geometric corrections grounded in physical modelling [Prieur et al., 2024]. Beyond correction, phenology-aware methods and self-supervised representation learning (e.g. Barlow Twins; Zbontar et al. 2021, Prieur et al. 2025) applied to multi-temporal data can learn encodings invariant to nuisance variation and domain shift yet sensitive to phenological identity, and with modest local fine-tuning could support scalable biodiversity pipelines. Progress ultimately requires integrating trait-informed spectral interpretation, sensor harmonisation and noise modelling, and temporal-aware machine learning.

**Data priorities.** Progress depends on well-labelled hyperspectral datasets spanning multiple seasons, years and sites, ideally with overlapping taxa to test generalisation (cf. NEON in the North American context; Marconi et al. 2022). Paracou has recently undergone a second hyperspectral scan which will be further complemented by equivalent acquisitions at Nouragues (French Guiana), providing a rare opportunity to quantify spatial and temporal transferability under realistic conditions and to identify which spectral features or representations remain stable across environmental and technical variability.

**Implications for hyperspectral sensing from space.** Our findings bear directly on the use of multispectral (e.g. Sentinel-2) and hyperspectral (e.g. EnMAP; forthcoming CHIME, HYPXIM) satellites to map forest composition, traits, and phenological diversity at scale. The most informative window, the FRE (748–775 nm), falls between Sentinel-2’s red-edge bands (B6 = 740 nm; B7 = 783 nm), and even with ideal placement its broad bandwidths (∼15– 20 nm) would smear the subtle differences critical for species discrimination; Sentinel-2’s ∼10 m pixels can capture large crowns, but its spectral granularity is insufficient. Current and planned hyperspectral missions offer the narrow, contiguous bands needed for FRE-scale cues, but their coarser pixels (≥30 m) under-resolve individual tropical crowns. Without robust unmixing methods to disentangle sub-pixel species mixtures, this spectral–spatial trade-off is a first-order bottleneck, compounded by atmospheric, view-geometry, phenological and illumination variability that satellite data amplify. Consequently, while trait or community-level patterns can be detected [Fassnacht et al., 2016, Féret and Asner, 2013, Aguirre-Gutiérrez et al., 2025], fine-grained species classification remains elusive under current capabilities. Multi-sensor fusion offers a partial route forward: because adjacent bands carry partially redundant information, broader-band satellite data may be usable with appropriate feature engineering, and field-validated airborne models co-registered with temporally matched satellite observations could support hybrid classifiers that bridge scales. For now, high-resolution airborne platforms remain essential both for advancing species-level mapping and for establishing the benchmarks that will guide next-generation satellite systems.

## 5 Conclusions

This study demonstrates that accurate, species-level mapping of tropical forest canopy trees at landscape scales is achievable by combining temporally fused CNN-based crown segmentation with high-resolution airborne hyperspectral imagery. At Paracou, consensus mapping across ten UAV RGB surveys improved segmentation F1 from 0.68 (single date) to 0.78 (ten dates), covering 86% of the test region’s canopy area, and LDA on 378 hyperspectral bands achieved a weighted F1 of 0.75 across 169 species, with repeated cross-validation confirming that on average 50 species (95% CI: 41–63) attain F1 ≥ 0.7 per fold and 15 do so reliably. Combining both steps, approximately 70% of the landscape’s canopy area was correctly mapped to species. Performance was nonetheless strongly skewed toward common species (macro-average F1 = 0.49 on the held-out test set; 0.48 across cross-validation folds), and many rare species with limited training data remained unclassifiable. The far-red edge (748–775 nm) was the most informative spectral region, and ablation confirmed that a reduced band set centred on it retains most classification power. These results advance substantially on previous studies limited to 20 or fewer species, but the findings are specific to a single, well-instrumented site with a decade of field data investment, accuracy depends critically on training data availability, and generalisation to other forests, sensors, or scales remains to be demonstrated. Progress toward operational species mapping will require well-labelled datasets across multiple sites and seasons, together with advances in transfer learning and sensor harmonisation.

## Supporting information

Supplementary Confusion Matrix

## Acknowledgements

J.G.C.B. was supported by the NERC C-CLEAR doctoral training programme (PDAG/501) and the Franklinia Foundation. S.J. was funded by a charitable donation from John Bernstein. The work (including access to high performance computing clusters) was supported by the Cambridge Centre for Earth Observation. Data collection in French Guiana was supported by CNES who funded the 2016 hyperspectral data over Paracou and Labex CEBA (ANR-10-LABX-25) who funded the UAV RGB collections and the field validation of manual crown segmentations as part of the Phenobs project. Thanks to Jean-Louis Smock (IRD), Ilona Clocher (CNRS), Isabelle Maréchaux (INRA), Chantal Geniez (IRD), Julien Engel (IRD), Tom Hattermann (CNRS), Géraldine Derroire (CIRAD), Patrick Heuret (INRA), and all staff at Paracou Research Station for help in the field. Thanks to Philippe Verley (IRD) for support in data processing.

## CRediT authorship contribution statement

JGCB wrote the manuscript and all authors contributed to the final version. JGCB, SJ, GV and DAC conceived the study design. JGCB and SJ ran the model training and supporting experiments. AL and CP developed the hyperspectral data from raw to analysable states. GV supervised the fieldwork and HS data collection. NB supervised the UAV data collection. TDJ contributed to methodological development and data interpretation. AM supervised the computational infrastructure.

## Use of AI tools

AI-based language tools (Claude, Anthropic) were used during manuscript preparation to improve the clarity and flow of text that had already been drafted by the authors. AI was not used to generate scientific content, interpret results, or design analyses; all intellectual contributions remain those of the named authors.

## Data availability

Upon acceptance the following will be archived with persistent identifiers. The field-validated crown polygons, with species labels, confidence scores, temporal validity dates and liana annotations, will be deposited on Zenodo, with schema and metadata described at umr-amap/ParacouTrees; precise trunk locations are subject to data-sharing agreements with the Paracou Field Station, but spatially masked versions will be available without restriction. Hyperspectral flight-line reflectance and UAV RGB orthomosaics will also be deposited on Zenodo, and the lidar-derived CHM is available through the Paracou data portal. All classification, cross-validation, ablation and consensus-fusion scripts, with environment files and model configurations, are at sadiqj/hyperspectral-nns and sadiqj/hyper-analysis; *detectree2* is at PatBall1/detectree2, with the pre-trained base model on Zenodo (doi:10.5281/zenodo.10522461).

## Footnotes

1 https://zenodo.org/records/10522461

2 https://scikit-learn.org/stable/modules/svm.html#complexity

