## Supplementary Materials for "Advances and limits of species-level canopy mapping in a hyperdiverse tropical forest: multi-temporal crown segmentation and airborne imaging spectroscopy"

#### S1 Three-way human manual segmentation comparison

This experiment helps to put into perspective the skill that machine learning algorithms can demonstrate in delineating tree crowns when human interpreters are taken as the ground truth.

To test how well different human interpreters agree when identifying tree crowns from imagery provided to them, two plots at Paracou were selected to be delineated by three expert human analysts. Three expert human analysts (familiar with remote sensing data and segmentation methods), Analysts A, B and C were asked to segment the tree crowns of two plots. The segmentation was performed in QGIS with the following data layers available: 2015 and 2016 RGB images and lidar CHMs, 2016 hyperspectral imagery (all spectral bands and PCA projected bands).

The three sets of delineations for each of the two plots were compared against each other to determine the degree of congruence. A match was granted when polygons from one set had a Jaccard/IoU $>$ 0.5 with one from another set.

**Plot 1 results**

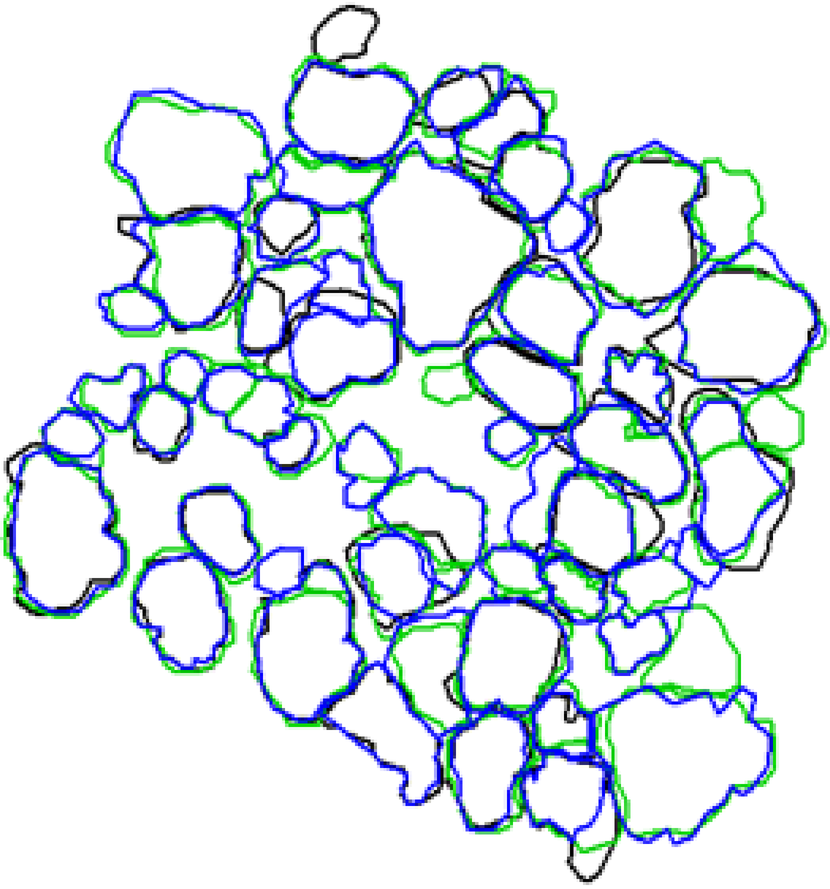

**Figure S1:** A comparison of manual human tree crown delineations (Plot 1). Set A: black, Set B: green, Set C: blue.

Comparison A-B: 28 congruent segments (Jaccard>0.5): 90% of A's crowns and 56% of B's.

Comparison A-C: 30 congruent segments (Jaccard>0.5): 97% of A's crowns and 60% of C's.

Comparison B-C: 42 congruent segments (Jaccard>0.5): 84% of B's crowns and 84% of C's.

**Plot 2 results**

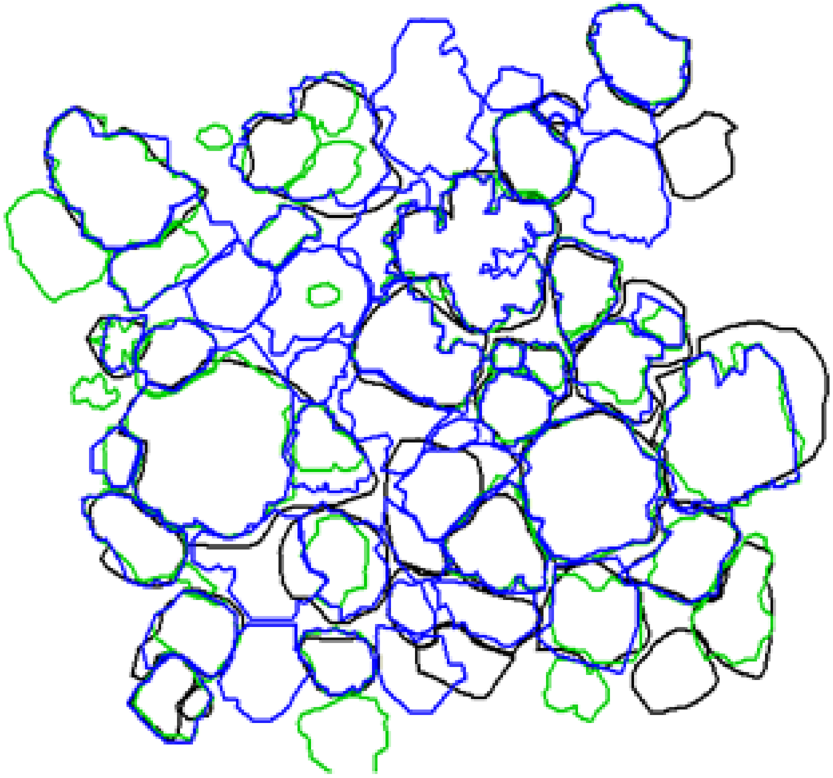

**Figure S2:** A comparison of manual human tree crown delineations (Plot 2). Set A: black, Set B: green, Set C: blue.

Comparison A-B: 19 congruent segments (Jaccard>0.5): 65% of A's crowns and 48% of B's.

Comparison A-C: 22 congruent segments (Jaccard>0.5): 76% of A's crowns and 46% of C's.

Comparison B-C: 28 congruent segments (Jaccard>0.5): 72% of B's crowns and 58% of C's.

The results show that humans can interpret images quite differently highlighting the challenge of achieving high accuracy with automated methods of tree crown delineation.

### S2 Remote sensing processing details

#### S2.1 RGB preprocessing

The RGB orthomosaics were compiled from the raw geotagged UAV photographs using structure from motion (SfM) photogrammetry in AgiSoft Metashape. The software aligns overlapping images to produce a sparse point cloud, refines it into a dense point cloud, and subsequently constructs a 3D mesh. This mesh, integrated with original photo textures, facilitates the creation of a Digital Elevation Model (DEM). The DEM, combined with the aligned images, allows for the generation of an orthomosaic, a georeferenced image free from perspective distortions. Supplying images across several dates in single blocks to the first steps of the SfM processing improves spatio-temporal coherency (Feurer and Vinatier 2018). Following this approach, instead of processing each date separately, five date blocks were supplied for the alignment and initial sparse point cloud formation establishing a common geometry between dates. The dates were then separated for the dense matching steps and final orthomosaic generation. We used an earlier airborne lidar dataset to assist in the positioning and alignment of the orthomosaics, providing a baseline layer for integrating and interpreting other remote sensing data.

##### S2.2 Hyperspectral preprocessing

Hyperspectral preprocessing is described in detail by Laybros et al. [2019, 2020]. A King Air B200 airplane flew at an average altitude of 920 m above ground level on 19-Sept-2016 (equivalent to approximately 900 m above the forest canopy, as reported in Table 1). Hyspex VNIR-1600 and Hyspex SWIR-384 (covering the 416-2500 nm wavelength range) were mounted to an aircraft side-by-side. The plane made 23 overpasses of the study site moving (North to South and South to North on consecutive overpasses) resulting in 23 separate, overlapping (~50%) flight lines of data. To merge the data from the two hyperspectral sensors without degrading the spatial resolution of the VNIR imagery, we resampled the SWIR imagery to 1 m using nearest-neighbour interpolation. Images were orthorectified and georeferenced at 1 m spatial resolution with the PARGE software using a canopy Digital Surface Model (DSM) produced from a lidar point cloud acquired over Paracou contemporaneously with the 2016 hyperspectral campaign [Laybros et al., 2019, 2020]. This acquisition is distinct from the November 2019 lidar scan listed in Table 1 and was used only for orthorectification, ensuring that the surface model represents the canopy as it stood at the time of the hyperspectral flight; the 2019 scan supplies the CHM used as the common spatial reference for crown delineation and co-registration (Section S2.3). Bands in the SWIR with a low signal to noise ratio due to water absorption peaks were removed leaving 378 of the 448 total bands. Per pixel illumination was calculated using the shadow detection method of Schläpfer et al. [2018]. Spectral information used to train and make predictions with the species classifiers was extracted from the overlapping flight lines rather than from a mosaic. This allows for valuable information to be retained as multiple views of individual crowns within the overlapping flight lines which has been shown to improve the classification performance. Spatial filtering decreases the local noise on each pixel and may improve the separability of objects in hyperspectral data. A spatial filter (mean of a $3 \times 3$ moving window) was applied to the flight line acquisitions. Brightness normalisation was then applied to each pixel spectrum: the reflectance value of each band was divided by the sum of that pixel's reflectance values across all bands, which has been shown to improve tree species classification [Dalponte et al., 2014]. Some machine learning classifiers are sensitive to the scale in which each feature (band in this case) is supplied with features that have a higher absolute variability tending to dominate. To address this, we applied the 'standard' scaling approach which standardises features by removing the mean (centring on zero) and scaling to unit variance.

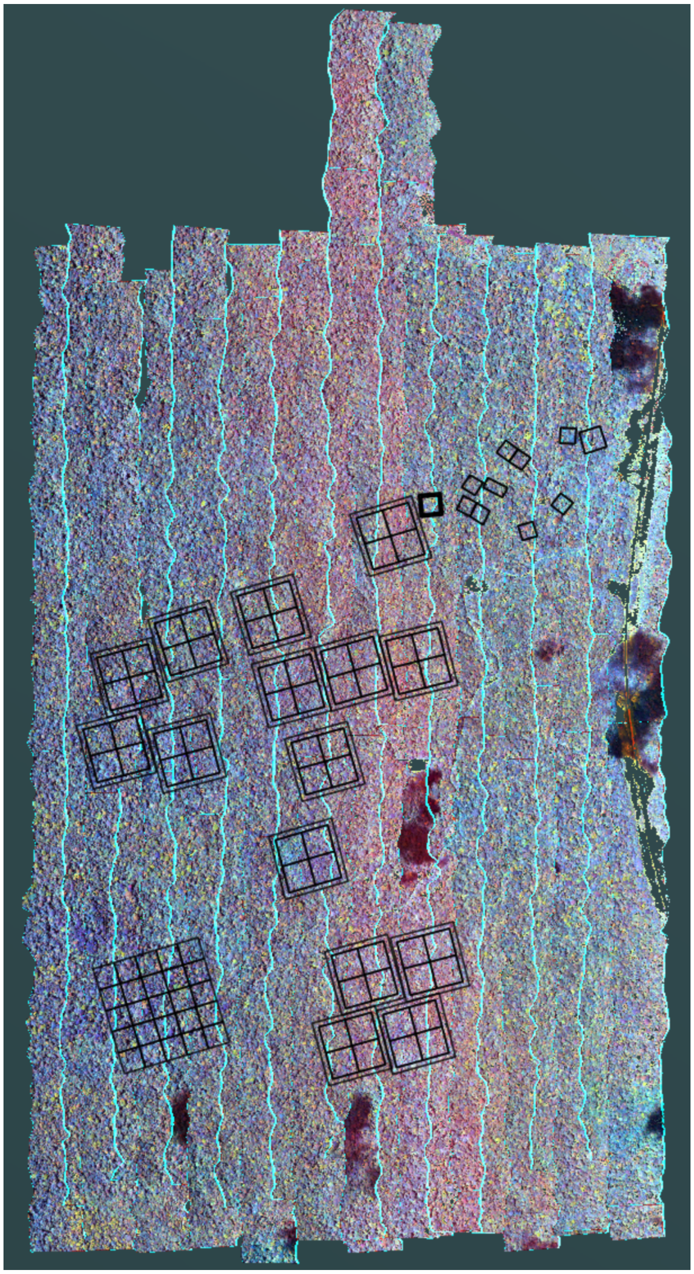

**Figure S3:** Hyperspectral flightlines over Paracou, French Guiana on 19-Sept-2016.

##### S2.3 Co-registration

Accurate co-registration of data from RGB and hyperspectral imagery was important to ensure spatial alignment. We used the Canopy Height Model (CHM) derived from the November 2019 lidar scan (Table 1) as the baseline layer, with all other data registered against it. This choice was due to the CHM's stability and precision in representing the physical landscape, providing a solid reference for co-registration. Eight control points were manually assigned across the different datasets, using identifiable features within the lidar CHM, such as the flux tower, roads and dominant trees; affine transformations were applied based on these. This co-registration process ensured that the crowns represented across the datasets corresponded to the same geographical location, serving as the foundation for subsequent analysis steps, including tree crown delineation and species classification. Although formal residual errors were not recorded during this manual process, visual inspection of the aligned datasets against the lidar CHM showed sub-pixel consistency for the identifiable features used as control points (flux tower, roads, emergent crowns). Given the 1 m resolution of the hyperspectral imagery and the 0.5 m CHM, residual misalignment is expected to be below 1 m for most of the study area, though localised errors may be larger in areas far from control points.

##### S2.4 Temporal compatibility of data sources

The remote sensing datasets span different acquisition dates (Table 1): UAV RGB imagery was collected over 2020–2021, hyperspectral imagery in September 2016, and the lidar CHM used throughout the analysis in November 2019. The field-derived crown database was built and curated between 2015 and 2023. Several features of the data and workflow accommodate this spread.

Two lidar acquisitions are involved and serve distinct purposes. An earlier scan, contemporaneous with the 2016 hyperspectral campaign, supplied the canopy surface model used to orthorectify the hyperspectral flight lines, so that orthorectification is epoch-matched to the imagery it corrects (Section S2.2). The November 2019 scan provides the CHM used as the common spatial reference for crown delineation and co-registration (Section S2.3), giving a geometrically stable reference frame against which all other layers are registered.

Each crown polygon in the database carries validity dates (`StartDate`, `EndDate`) that record when mortality, major branch fall, or significant crown change was observed, allowing crowns to be filtered to match the epoch of each sensor (Section S3). Of the 3,500 crowns in the database, 3,256 had valid species assignments and sufficient temporal overlap with the 2016 hyperspectral acquisition for inclusion in the classification task; crowns whose canopy had changed substantially between 2016 and the date of delineation or field verification were excluded or updated.

Crown growth in this forest is also slow relative to the inter-sensor time gaps: undisturbed crowns do not change shape substantially over the 3–5 year intervals between acquisitions (Section S3.1), and the principal sources of change – mortality and branch fall – are discrete events that the database explicitly tracks. UAV-based segmentation training and evaluation therefore used only crowns valid during the 2020–2021 RGB campaign, while species classification training and evaluation used crowns valid during the 2016 hyperspectral acquisition. Segmentation and classification are thus trained and evaluated on separate, epoch-matched crown sets, so each stage is assessed against imagery that matches the canopy state it represents.

#### S3 Tree crown dataset

The dataset and data description is available in this GitHub repository: https://github.com/umr-amap/ParacouTrees.

To train machine learning algorithms and evaluate automatic tree crown delineation and species identification from remote sensing data it is necessary to have an extensive ground truth map of tree crowns. Generating this takes time and attention to the specific attributes of each crown. Careful ground validation is necessary to have confidence in correct individual/species assignment and delineation (avoiding over/under segmentation).

##### S3.1 Premises of the dataset

The growth of crowns is relatively slow, meaning an undisturbed crown will not change its shape significantly between scans/field missions. Creating crown polygons is a time consuming process as it requires a careful comparison/contemplation of the different modalities (and time steps thereof) of scans against field inventory data.

Significant changes to the crown are due to:

- Tree death
- Branch fall

It is not feasible or time efficient to produce a new set of crowns for each new scan. Instead, the crowns are updated (by hand) when a significant change is detected. The fields StartDate and EndDate are used to track the validity of a crown. fid_1 is a unique identifier for an individual tree - multiple, temporally distinct, crown polygons may be associated with an fid_1. StartDate and EndDate are set to NULL when a crown is created. If a crown is seen to no longer be valid (e.g. due to a branch fall or mortality), EndDate is set to the date of the scan that shows this. A new crown may be created if an existing crown changes due to branch fall, if a significant portion of an existing crown is revealed by a branch fall or the mortality of an occluding tree, or if a new tree is discovered. Full details of the fields are given below.

##### S3.2 Fields of dataset

A series of fields are used to describe the crown polygons:

- fid (int): unique identifier for each crown polygon
- fid_1 (int): a unique identifier for individual trees (not polygons). This can be useful to track individuals if a crown has changed significantly through time (see StartDate, EndDate).
- Site (str): Location of data collection (e.g. Paracou, Nouragues etc.)
- PlotOrg (str): Necessary at Paracou (CIRAD, CNES or INRA). This helps in linking the polygons to the inventory datasets.
- PlotNum (int): plot number
- SubPlot (int): some plots have subplots contained within them
- LocalID (int): the tree number as recorded on the tree's tag

- TrunkMatch (int): 1,2,3,4 These integers describe how well the crown polygon (as delineated from the remote sensing data) has been matched to a trunk in the field.
- CrownIntegrity (int): 1,2,3,4 These integers describe how sure we are that a delineated polygon is that of a single, complete crown.
- Lianas (bool): as to whether lianas are present in the crown of the tree delineated
- StartDate (date): Date at which the crown becomes visible or has changed shape
- EndDate (date): Date at which the crown becomes absent or has changed shape
- Dead (bool): a crown might be present or belong to a dead tree
- GroundValid (bool): has the crown been checked in the field?
- Creator (str): name of the person to have made the polygon
- Comments (str): for any comments before or in the field
- BaseLayer (str): which remote sensing data source has been used as the "anchored" location of the crown

To be included in the tree species classification training and testing a crown had to satisfy CrownIntegrity $\leq$ 2 & TrunkMatch $\leq$ 2 & GroundValid == TRUE.

### S4 Multi-date tree crown fusion (consensus mapping)

#### S4.1 Cross-date matching and grouping

We concatenated all per-date polygon prediction sets into a single pool and identified cross-date matches using the Intersection-over-Union (IoU) metric:

$$\mathrm{IoU}(A,B) = \frac{\mathrm{area}(A \cap B)}{\mathrm{area}(A \cup B)}$$

A **significant match** was defined as **IoU $\geq$ 0.75**. For each polygon, all other polygons in the pool were scanned to find such matches; polygons linked by significant matches were grouped as candidate observations of the **same tree crown** across dates. In a 10-date stack, any polygon can therefore have **0–9** matches (i.e., be confirmed on up to nine other dates). A group size of one (no matches) was retained and processed identically in the steps below.

#### S4.2 Confidence-weighted boundary fusion

For each matched group, we produced a single consensus polygon via boundary normalisation and confidence-weighted averaging:

1. **Boundary normalisation.** Each member polygon was resampled to 300 boundary vertices.

2. **Vertex alignment.** For each polygon, the starting vertex was the boundary point with minimum $x$+$y$; vertices were then ordered clockwise so that vertex index j ∈ 1,. . . ,300 corresponds across polygons.

3. **Confidence-weighted averaging.** Let $x_{ij}, y_{ij}$ be the *j*-th vertex of the *i*-th polygon in the group, and let $w_i \in [0, 1]$ be its per-polygon confidence. The fused vertex is

$$\bar{x}_j = \frac{\sum_i w_i x_{ij}}{\sum_i w_i}, \quad \bar{y}_j = \frac{\sum_i w_i y_{ij}}{\sum_i w_i}$$

This yields an averaged boundary $\{(\bar{x}_j, \bar{y}_j)\}_{j=1}^{300}$.

4. **Area and centroid adjustment.** The fused polygon was scaled so that its area equals the mean area of the original polygons, with scaling performed about the mean centroid of the originals.

We recorded two attributes for each fused polygon:

- **Summed confidence** $C_{\text{sum}} = \sum_i w_i$ which aggregates member confidences and implicitly reflects cross-date agreement;
- **Combination count** n, the number of polygons fused (equivalently, the number of dates on which the crown was delineated).

##### S4.3 Landscape-level filtering and space-filling selection

We constructed the final spatial mosaic of consensus crowns via a greedy, confidence-ordered selection:

1. **Quality filter.** Discard any fused polygon failing the minimum confidence threshold (see Section S4.4).
2. **Greedy placement.** Sort remaining polygons by **descending** Csum. Iteratively place polygons into the landscape provided they do **not** conflict with already placed polygons, where a conflict is defined by **IoU** $\geq$ **0.2** with any placed polygon. (Equivalently, we only accept candidates whose overlap with all placed polygons satisfies **IoU** $<$ **0.2**, enforcing *minimal overlap*.)
3. **Result.** The accepted set provides a **spatio-temporal integration** of tree-crown predictions, with each polygon representing the average **location and outline** of a crown across multiple time points and carrying agreement-aware confidence.

##### S4.4 Parameter settings and tuning

All thresholds and settings were tuned **only on training crowns** and then **fixed** for evaluation in the held-out test regions:

- Cross-date match threshold: **IoU** $\geq$ **0.75** (S.4.1).
- Boundary normalisation: **300 vertices** per polygon; start at **min(x+y)** (S.4.2).
- Confidence aggregation: **sum of member confidences** Csum and **combination count** n (S.4.2).

- Quality filter: Csum>0.1 × n (S.4.3).
- Space-filling conflict threshold: **IoU < 0.2** required for acceptance (i.e., conflicts defined as **IoU ≥ 0.2**) (S.4.3).

##### S4.5 Notes on computational procedure

- Matching is performed on the union of per-date predictions; grouping uses transitive closure over pairs with **IoU ≥ 0.75**.
- The confidence-weighted fusion preserves sharper, higher-confidence boundaries while damping date-specific artefacts.
- The greedy selection produces a **non-overlapping** (up to IoU < 0.2) set of crowns prioritised by cross-date agreement and per-date confidence.

Code implementing the full pipeline is available on GitHub[3].

#### S5 Species classification: overall accuracy

##### S5.1 Tuned model parameters

Model tuning was based on pixelwise k-fold cross validation with the constraint that pixels from any individual crown must be contained within a single fold (i.e. not split between training and validation folds) and a grid search of hyperparameters. The tuned model parameters are presented in Table S1.

##### S5.2 Exploratory analyses and notes on model selection and training

A range of other modelling techniques were tried but not included in the final model testing/selection as they failed to be competitive in terms of species prediction and were often highly computationally intensive. A judgement to drop these was made as it became clear that they would not help to achieve the aims of the study and could draw resources away from other necessary components of the work.

The methods can be split into modelling architectures/pipelines and augmentation/sampling/weighting techniques. The modelling approaches for which we did preliminary testing but did not take further, due to poor relative performance:

- **UMAP**[6] is a non-linear dimension reduction technique that we thought would be effective for representing the hyperspectral data. We tried the unsupervised and supervised implementation to generate potential features for classification but despite a broad sweep of hyperparameters we could not create an effective model.

[3] `https://github.com/PatBall1/detectree2/blob/f996564bfcbaed1ff0ef13a63ea3e62f47252731/detectree2/models/outputs.py#L439C10-L439C10`

[6] https://umap-learn.readthedocs.io/

**Table S1:** Tuned model (hyper)parameters for the best performing models. The range of parameters trialled during tuning are given in brackets after the optimised value. Aside from the MLP which was implemented in PyTorch, all models were implemented in scikit-learn*[4]

| Model | Best parameters [trial range] | Notes |
|---|---|---|
| MLP | - Four-layer fully-connected MLP - Two [2, 10] hidden layers of dimension 128 [16, 512] - Rectified Linear Units (ReLU) - Dropout of 0.05 [0.01, 0.4] - Learning rate of 3e-4 [1e-5, 0.01] - Cosine annealing over 30 epochs - AdamW optimiser - Label smoothing of 0.2 [0.05,0.3] | We also experimented with various model architectures, including CNNs, and data augmentation techniques like MixUp and SMOTE; however, these approaches yielded inferior performance. |
| KNN | Nearest neighbours: 64 [2, 512] | |
| Logistic Regression | C = 10 [0.01, 1000] | |
| QDA | reg = 0.1 [0.005, 0.5] | |
| SVM | C = 1 [0.001, 100] kernel = linear ['linear', 'poly', 'rbf', 'sigmoid'] degree [2, 4] | Linear kernel was best performing – knowing this training times could be improved with LinearSVC in the liblinear implementation[5] |
| RandomForest | n_estimators=100 [50, 120] criterion='gini' No max_depth [5, None] min_samples_split=2 [2, 10] max_features = sqrt ['none', 'sqrt', 'log2'] min_samples_leaf=1 [1, 4] | |
| LDA | solver = svd ['svd', 'lsqr', 'eigen'] | |

- We designed a **1D CNN** that convolved along the spectral dimension of the pixels again. We hoped that it could effectively represent the spectral features of the data. Again, despite an initial broad sweep of hyperparameters we could not find an effective model.

- We also tried a model pipeline that first performed a dimensionality reduction (using PCA, LDA and UMAP) on the hyperspectral data to produce a reduced set of features that were then fed into a secondary classifier such as an SVM. However, these combinations did not show any advantage over the classifiers used with all available features.

We tried a few other techniques to help boost the accuracy of the classifications, especially of the less well represented classes:

- **Synthetic Minority Oversampling Technique (SMOTE)**[7] is an augmentation technique for boosting the representation of less well represented classes. We applied this to the training pixels but in our cross-validation, while we observed a marginal boost to some less well represented classes, there was a drastic reduction in overall accuracy.

- Setting class_weight="balanced" in scikit-learn sets the class weights to be inversely proportional to their frequency, preventing the model from becoming biased toward the majority class. With this applied, we also observed a marginal boost to some less well represented classes that we did not judge to be worth the exchange for a drastic reduction in overall accuracy.

[7] https://arxiv.org/abs/1106.1813

- **MixUp**[8] is a data augmentation technique that creates new training examples by linearly interpolating between pairs of original training samples and their corresponding labels. We thought that this would improve our models' generalisation ability and robustness, but it yielded inferior results in comparison to using the non-augmented data.

These methods did not help us to achieve our aim of accurate landscape scale classification, so they were not included in the final training approaches.

We additionally tested several classifier families not included in the main comparison (Table 3) but frequently recommended in the hyperspectral species classification literature. HistGradientBoostingClassifier and PLS Discriminant Analysis (PLS-DA) – a supervised dimensionality-reduction method designed for correlated spectral data [Richter et al., 2016] – both scored below 0.30 (Table S2). Elastic-net logistic regression (L1/L2 penalty via SGD) performed similarly (0.26), confirming that L1-induced band sparsity does not help when the discriminant signal is distributed across many correlated bands. LDA with Ledoit-Wolf covariance shrinkage [Ledoit and Wolf, 2004] scored 0.67, slightly *below* standard LDA (0.73), indicating that LDA's SVD solver already provides effective implicit regularisation for this dataset. GaussianNB (0.06) served as a diagnostic control: its assumption of independent features (diagonal covariance) is catastrophically wrong for correlated spectral bands, confirming that inter-band covariance modelling is essential. Gradient Boosting, XGBoost, LightGBM, and Extra Trees were also tested during exploratory work (hyperparameter ranges in Table S3) but failed to converge to competitive accuracy within reasonable compute budgets, consistent with the Random Forest result (0.44) and the broader finding that tree-based ensembles lack the implicit dimensionality reduction that makes LDA effective on high-dimensional correlated spectra.

##### S5.3 Relationship between overall accuracy and the number of species represented in the training pool

To examine how classification accuracy varies with the number of classes (species) present, we reran training and testing with LDA (the best performing model) on a series of artificially restricted datasets. Starting from the full dataset, the individuals of the least well represented species were iteratively removed from the dataset until just two species remained. We checked how classification accuracy varied when: (i) the testing pool of species was kept to the complete set with 169 species while the training pool was restricted; (ii) the testing pool was restricted in line with the reduction in the training pool.

Accuracy with the "Complete" test pool decreased as species were removed from the training data whereas the accuracy of the "Matched to train" test pool increased (Fig. S4). There was no penalty to overall (landscape) accuracy for including as many species as possible. This demonstrated the value of attempting to include all represented species in the model training and classification.

[8] https://arxiv.org/abs/1710.09412

**Table S2:** Sample of the exploratory results performed with cross validation on the training data with stratified grouping based on individual crowns (pixels from within the same crown could not be in both training and validation folds). The parameters from the grid search that gave the highest mean CV accuracy were recorded. "unnorm" runs were performed on the unnormalised spectral data. Fit times were not recorded for all sweeps.

| Model | Best parameters | Mean CV accuracy | Std. dev. | Fit Time (s) |
|---|---|---|---|---|
| LDA | | 0.71 | 0.02 | |
| KNN | 'n_neighbors': 64 | 0.37 | 0.01 | |
| Logistic Regression | 'C': 10 | 0.55 | 0.02 | |
| SVM | 'C': 1, 'kernel': 'linear' | 0.63 | 0.02 | |
| QDA | 'reg_param': 0.1 | 0.50 | 0.02 | |
| KNN | 'n_neighbors': 128 | 0.36 | 0.01 | |
| Logistic Regression | 'C': 80 | 0.55 | 0.02 | |
| PCA_SVM | 'pca__n_components': 50, 'svm__C': 0.1, 'svm__kernel': 'linear' | 0.57 | 0.02 | |
| LDA_SVM | 'svm__C': 1, 'svm__kernel': 'rbf' | 0.69 | 0.02 | |
| LDA | 'n_components': None, 'shrinkage': None, 'solver': 'svd' | 0.71 | 0.02 | 312.34 |
| LDA_unnorm | | 0.70 | 0.02 | 99.39 |
| SVM | 'C': 1, 'degree': 2, 'gamma': 'scale', 'kernel': 'linear' | 0.63 | 0.02 | |
| QDA_unnorm | 'reg_param': 0.05 | 0.51 | 0.01 | 12.83 |
| Random Forest_unnorm | | 0.40 | 0.01 | 63.48 |
| QDA_unnorm | 'reg_param': 0.00315 | 0.56 | 0.01 | 11.82 |
| LDA_unnorm | 'shrinkage': None, 'solver': 'lsqr' | 0.70 | 0.02 | 13.49 |
| LDA | 'shrinkage': None, 'solver': 'svd' | 0.71 | 0.02 | 27.27 |
| QDA | 'reg_param': 0.0045, 'tol': 1e-07 | 0.56 | 0.02 | 13.61 |
| LDA | 'shrinkage': 7.4e-05, 'solver': 'lsqr', 'tol': 1e-07 | 0.71 | 0.02 | 14.80 |
| SVM | 'C': 0.5, 'gamma': 'scale', 'kernel': 'linear' | 0.63 | 0.02 | 6596.78 |
| SVM | 'C': 0.4, 'gamma': 'scale', 'kernel': 'linear' | 0.63 | 0.02 | 8607.83 |
| SVM | 'C': 0.43, 'gamma': 'scale', 'kernel': 'linear' | 0.63 | 0.02 | 5852.08 |
| Random_Forest | | 0.44 | 0.03 | 83.77 |
| Random_Forest | 'n_estimators': 120 | 0.44 | 0.03 | 212.58 |
| Random_Forest | 'n_estimators': 250 | 0.44 | 0.03 | 511.01 |
| Random_Forest | 'max_depth': None, 'n_estimators': 250 | 0.44 | 0.03 | 784.57 |
| Random Forest | 'max_features': 'sqrt', 'n_estimators': 50 | 0.43 | 0.03 | 63.23 |
| *Extended comparison (378 bands, 169 species, crown-grouped 5-fold CV, weighted F1):* | | | | |
| HistGradientBoosting | 'max_iter': 200, 'max_depth': 6, 'lr': 0.1 | 0.29 | 0.02 | 59.72 |
| PLS-DA | 'n_components': 50 | 0.28 | 0.01 | 964.22 |
| Elastic-net Logistic (SGD) | 'l1_ratio': 0.5, 'alpha': 1e-4 | 0.26 | 0.01 | 407.34 |
| LDA (Ledoit-Wolf shrinkage) | 'solver': 'lsqr', 'shrinkage': 'auto' | 0.67 | 0.02 | 32.00 |
| GaussianNB | | 0.06 | 0.01 | 488.88 |

**Table S3:** Hyperparameter ranges for a sample of the exploratory analyses. Model types listed here but not listed in Table S2 failed to converge or took a prohibitively long time to train.

| Model | Hyperparameters |
|---|---|
| LDA | solver: ['svd', 'lsqr', 'eigen']; shrinkage: [None, 0.000074, 0.000077]; tol: [1e-7, 1e-6, 1e-5] |
| QDA | tol: [1e-7, 1e-6]; reg_param: [0.0047, 0.0043] |
| GaussianProcess | max_iter_predict: [100] |
| Random Forest (basic) | n_estimators: [50, 200, 300]; max_features: ['None', 'sqrt', 'log2'] |
| Random Forest (extended) | n_estimators: [50]; max_depth: [5, 10, 20, None]; min_samples_split: [2, 5, 10]; max_features: ['None', 'sqrt', 'log2']; min_samples_leaf: [1, 2, 4] |
| Random Forest (large) | n_estimators: [150, 200, 250, 300]; max_depth: [None]; min_samples_split: [2, 5, 10]; max_features: ['None', 'sqrt', 'log2']; min_samples_leaf: [1, 2, 4] |
| Gradient Boosting | n_estimators: [50, 100, 150]; learning_rate: [0.01, 0.1, 1]; max_depth: [5, 10, 20, None] |
| XGBoost | n_estimators: [50, 100, 150]; learning_rate: [0.01, 0.1, 1]; max_depth: [3, 5, 7] |
| LightGBM | n_estimators: [50, 100, 150]; learning_rate: [0.01, 0.1, 1]; max_depth: [3, 5, 7] |
| Extra Trees | n_estimators: [50, 100, 150]; max_depth: [3, 5, 7] |
| KNN | n_neighbors: [128, 256, 512] |
| Logistic Regression | C: [0.1, 1, 10, 100]; solver: ['liblinear', 'saga']; penalty: ['l1', 'l2', 'elasticnet', 'none'] |
| PCA_SVM | pca__n_components: [10, 20, 30, 50]; svm__C: [0.1, 1, 10]; svm__kernel: ['linear', 'rbf'] |

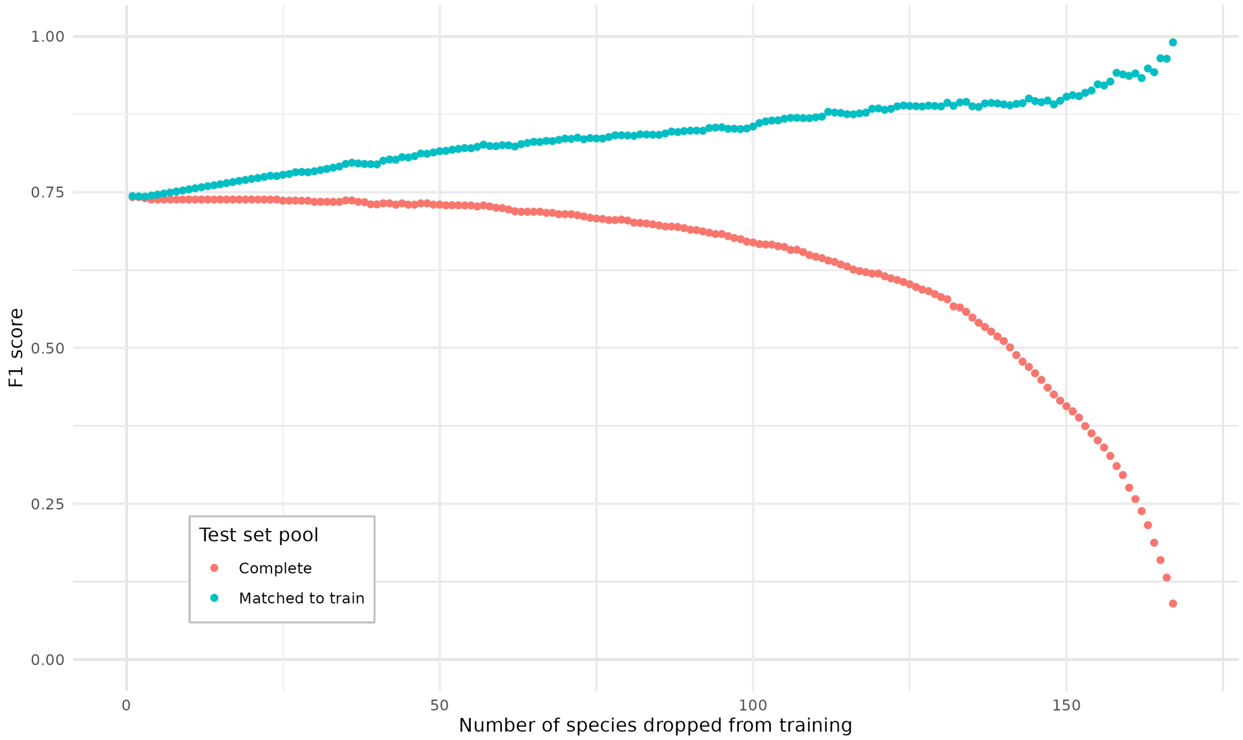


**Figure S4:** The relationship between weighted F1 of predictions on the test set and the number of species dropped from the training pool. The species were removed in order of least well represented to most well represented (with respect to number of individuals) starting with the full set and ending with just the two most well represented species. In the "Complete" set of experiments, the test set was maintained so that the full crown set of 169 species was tested. In the "Matched to train" set, the testing pool of species was reduced in line with the reduction in the training set.

### S6 Waveband importance

#### S6.1 Feature importance for LDA

In linear discriminant analysis (LDA), the discriminant axes are defined as linear combinations of the original features that maximise the ratio of between-class to within-class variance. Mathematically, this is formulated as a generalised eigenproblem,

$$\mathbf{S}_B\mathbf{w} = \lambda\mathbf{S}_W\mathbf{w}$$

where S_B and S_W are the between-class and within-class scatter matrices, respectively, and w is the eigenvector defining a discriminant axis. The raw coefficients of the discriminant function correspond to the elements of w, and directly specify how features are combined to form each discriminant score. Because raw coefficients are sensitive to the measurement units and variances of the predictors, they are not directly interpretable as feature importance.

To provide interpretable and comparable measures, we computed two complementary quantities.

**Variance-weighted squared scalings (standardised coefficients).** When the input features are standardised to unit variance prior to fitting (as in our pipeline), the raw scalings become standardised discriminant coefficients that are comparable across features. To obtain a single importance value per band aggregated across all K discriminant axes, we used the variance-weighted squared scalings method (Peterson and Mahajan, 1976):

$$\text{importance}_j = \textstyle\sum_k (w_{jk}^2 \cdot \lambda_k)$$

where w_jk is the standardised scaling of band j on axis k and lambda_k is the proportion of between-class variance explained by axis k. This aggregation is sign-invariant (via squaring) and gives greatest weight to the discriminant axes that contribute most to class separation. The resulting vector is normalised to sum to one. Because scalings measure the *unique* (partial) contribution of each band after controlling for all other bands, they can be sensitive to multicollinearity: when adjacent spectral bands are highly correlated, the model may concentrate its weight on a small number of bands that serve as representatives for a correlated group, inflating their apparent importance relative to their neighbours.

**Variance-weighted squared structure coefficients (loadings).** Structure coefficients are defined as the bivariate correlation between each original feature and the discriminant scores (Courville and Thompson, 2001):

$$r_{jk} = \text{cor}(x_j, Z_k)$$

where $x_j$ is the j-th predictor and $Z_k = Xw_k$ is the score on axis k. These capture the *total* association between each band and species discrimination, including information shared with correlated bands, and are robust to multicollinearity. We aggregated them in the same way: $\text{importance}_j = \sum_k (r_{jk}^2 \cdot \lambda_k)$, normalised to sum to one.

**Cross-validation stability.** To assess how robust the importance rankings are to the particular training sample, we extracted both metrics from LDA models trained in each of the 100 repeated cross-validation folds described in Section 2.7. For each method, we computed the mean and standard deviation of each band's importance across folds, as well as the mean pairwise Spearman rank correlation between all fold pairs (sampled at 500 pairs) as an overall stability metric.

**Comparison of methods.** The two metrics answer different questions and yield strikingly different importance profiles (Fig. S5). Variance-weighted squared scalings concentrate 91.5% $\pm$ 0.3% of total importance in the FRE (748–775 nm), reflecting that this narrow region captures the most non-redundant discriminative information. Structure coefficients, by contrast, spread importance evenly across the spectrum, with the FRE contributing only 2.5% $\pm$ 0.1%. This divergence is expected: with 378 highly correlated spectral bands, many bands carry species-relevant information (high structure coefficients), but this information is largely shared among adjacent bands. The scalings-based metric identifies where the classifier concentrates its discriminative power – the bands that carry information not available elsewhere. Both metrics showed high stability across CV folds (mean Spearman rho = 0.997 for scalings, 0.979 for structure coefficients), with the top-10 bands by scalings overlapping 95.4% on average across folds.

##### S6.2 Feature importance validation (ablation tests)

To further examine the importance of different spectral bands, we ran a series of ablation experiments. We used the average of the feature scalings at each band from the trained Linear Discriminant Analysis model to assign importance values and produce four different orderings of the spectral bands. For each ordering we then progressively removed bands (by setting them to zero) and then trained and tested a new LDA model. The four configurations were:

- **Ordered** in which we sorted the spectral bands by their respective feature scaling in descending order (the most "important" band according to its importance was removed first and the least important band removed last)
- **Reverse** used the same values as above but sorted in ascending order (the least important band was removed first and the most important band removed last)
- **Cluster shuffle** involved chunking the spectral bands into fifteen clusters and then performing a random shuffle
- **Random** randomly shuffled the spectral bands

The resulting test-set F1 scores for each configuration are shown in Fig. S6. Performance decreases quickest in the ordered configuration, followed by cluster shuffle, reverse and finally random. This indicates several things:

1. Feature scalings from LDA determine the bands necessary for discriminating species, as shown by the rapid decrease in F1 performance from the ordered configuration and conversely the slow decrease of the reverse configuration
2. There is a significant amount of redundancy across bands. In the random configuration more than half of the bands can be removed without significant performance degradation

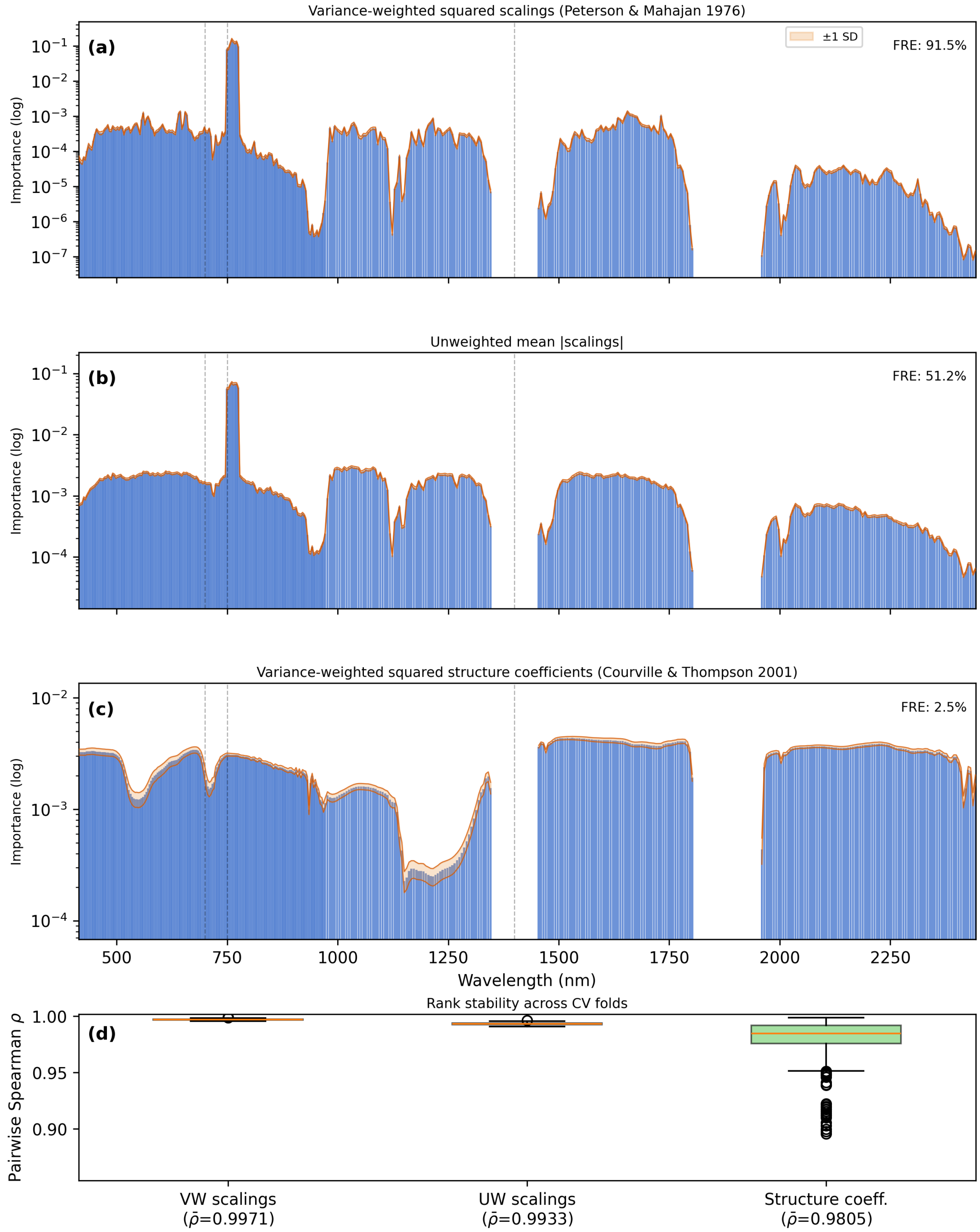


**Figure S5:** Comparison of three band importance metrics across 100 cross-validation folds. (a) Variance-weighted squared scalings (Peterson and Mahajan, 1976), which measure the unique contribution of each band; (b) unweighted mean absolute scalings; (c) variance-weighted squared structure coefficients (Courville and Thompson, 2001), which measure total association. Bars show the CV mean; shaded envelopes show $\pm$ 1 SD. The grey band highlights the far-red-edge (748–775 nm). (d) Distribution of pairwise Spearman rank correlations between fold importance vectors for each method, quantifying ranking stability.

3. There exists localised information which is necessary for discriminating species. This is evidenced by the decreased performance of the clustered shuffle configuration when compared to random

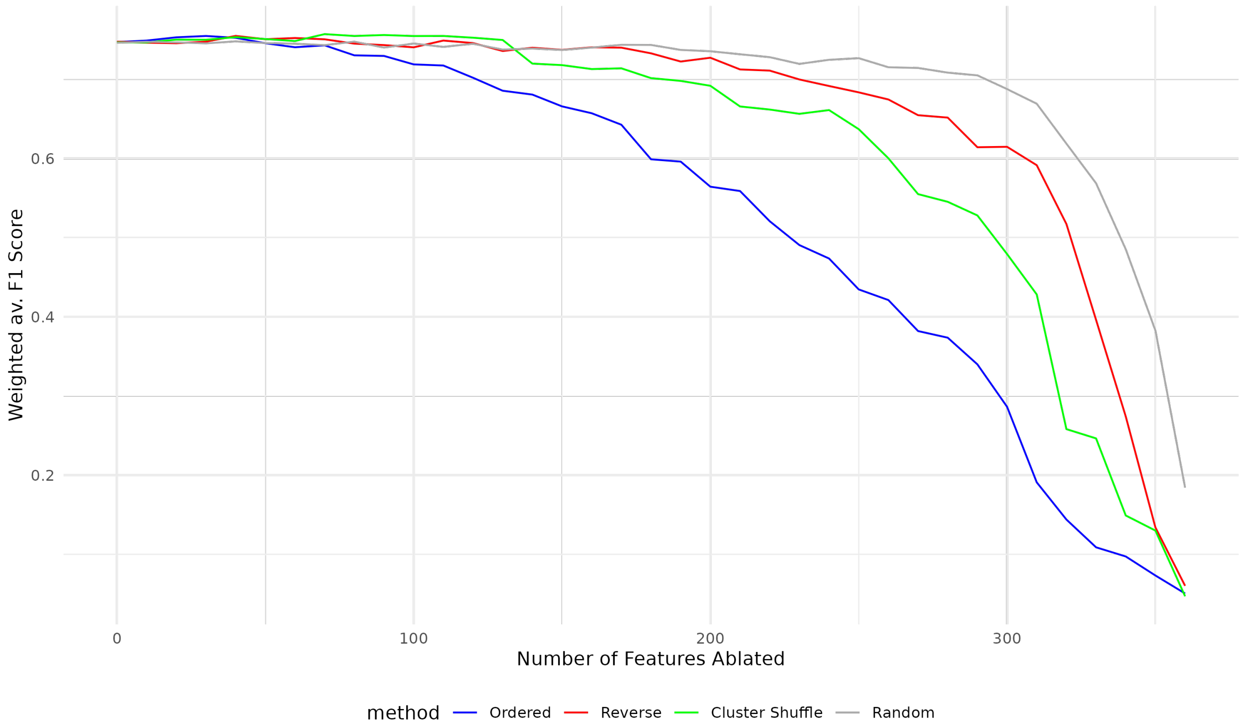


**Figure S6:** Weighted F1 of crown species classification predictions for the LDA classifier as bands are progressively ablated (i.e. standardised values set to 0 prior to training) in blocks of 10. The "Ranked" curve shows the effect of removing bands in order of feature importance starting with the most important bands. The "Reverse" curve shows the effect of removing bands in reverse order of importance. The "Cluster Shuffle" curve shows the effect of removing bands in an order which shuffled randomly 15 (k-means) clusters of band importance (retaining a realistic data structure as feature importance tends to cluster in specific regions). The "Random" curve shows the effect of removing bands after a random shuffle, which meant information across all spectral regions was available to the LDA for much longer than in the other methods.

##### S6.3 Feature selection and dimensionality reduction comparison

To directly test whether prior feature selection or dimensionality reduction could improve on full-band LDA, we conducted a systematic comparison using only model-agnostic feature selection criteria (avoiding circularity from using LDA-derived rankings to evaluate LDA). All experiments used stratified-group 5-fold cross-validation with crown-level majority-vote evaluation, matching the methodology of Section 2.7. For each value of $K$ in {10, 20, 50, 100, 150, 200, 300, 378}, we trained LDA on: (i) the top-$K$ bands ranked by mutual information (MI) with the species label [Kraskov et al., 2004], a filter-based criterion independent of any classifier; (ii) $K$ bands chosen uniformly at random (averaged over 10 random seeds as a null baseline); and (iii) $K$ principal components from PCA. No approach exceeded the full-band LDA baseline (weighted F1 = 0.727; Fig. S7). MI-based selection reached F1 = 0.70 by 200 bands but never surpassed full-band performance. PCA+LDA underperformed at every dimensionality tested (best: PCA(168)+LDA, F1 = 0.70), consistent with unsupervised projection discarding supervised discriminant structure.

Random selection was competitive at higher $K$ – reflecting the substantial spectral redundancy – but never exceeded the full-band baseline. MNF transformation was not tested separately as it is closely related to PCA and yields near-identical results for well-calibrated airborne hyperspectral data [Green et al., 1988]. These results confirm that LDA's implicit regularisation handles the full feature space effectively and that prior band selection or dimensionality reduction provides no net benefit for this classification task.

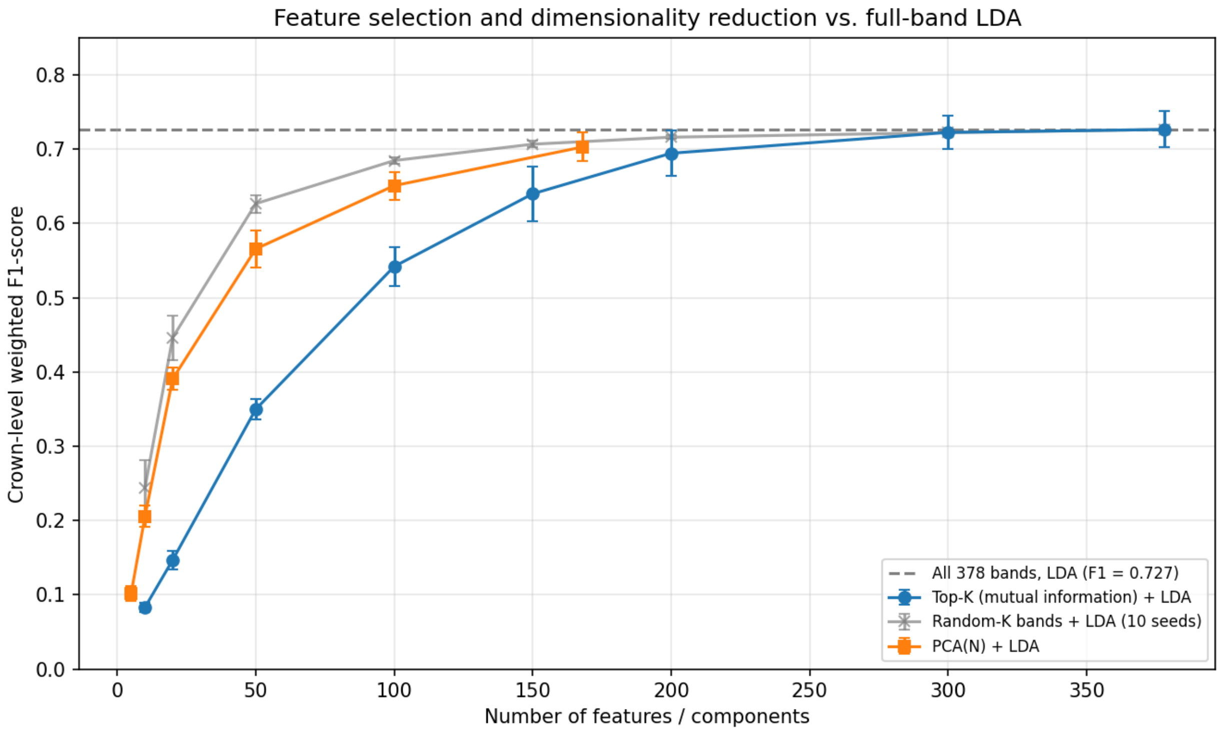


**Figure S7:** Crown-level weighted F1-score as a function of the number of retained features or principal components, comparing three model-agnostic feature selection strategies against full-band LDA (dashed line). Bands are ranked by mutual information with the species label (blue), or selected uniformly at random (grey; mean of 10 seeds). PCA dimensionality reduction (orange) serves as an unsupervised alternative. No approach exceeds the full-band LDA baseline (F1 = 0.727).

#### S6.4 Multi-classifier comparison across feature sets

Concerns that LDA's superior performance over Random Forest (RF) and Support Vector Machines (SVM) might arise from overfitting in high-dimensional feature space motivated a direct multi-classifier comparison across varying feature dimensionalities. We trained LDA, RF (100 trees, balanced class weights), and a linear SVM (SGD-based, balanced class weights) on the same MI-ranked Top-$K$ band subsets ($K \in \{10, 20, 50, 100, 200, 378\}$) and on a set of 15 standard vegetation indices (VIs). The VIs – NDVI, EVI, PRI, CRI1, ARI, MCARI, NDRE, $CI_{green}$, $CI_{RE}$, SIPI, $MSR_{RE}$, NDWI, SWIR ratio, red-edge position, and red-edge slope – span the major biochemical and structural traits used in hyperspectral vegetation studies [Fassnacht et al., 2016, Hennessy et al., 2020, Clark et al., 2005, Ferreira et al., 2023]. All feature sets are model-agnostic, ensuring a fair comparison.

LDA substantially outperformed both RF and SVM at every feature dimensionality (Fig. S8). Critically, the performance gap *widened* with increasing dimensionality – the opposite of what would be expected under an overfitting hypothesis. At full bandwidth (378 bands), LDA achieved F1 = 0.73, compared to RF at 0.33 and SVM at 0.26, while at $K = 10$ the three classifiers were comparably weak (F1 = 0.08, 0.13, 0.01 respectively). This pattern is consistent with LDA's linear projection into discriminant space – which implicitly regularises by reducing dimensionality to at most $K - 1$ axes – becoming increasingly effective as more spectral information becomes available, while RF and SVM lack this built-in dimensionality reduction and are more susceptible to the curse of dimensionality with crown-grouped cross-validation that prevents within-crown leakage.

On the 15 VIs, all classifiers performed poorly (LDA 0.26, RF 0.29, SVM 0.14), confirming that a small set of generic spectral indices – even those designed to capture the major biochemical axes of vegetation variation – cannot span the 168-dimensional discriminant space required to separate 169 species. This result is consistent with the finding that band selection studies achieving high accuracy typically involve far fewer species [e.g. 7–20 species; Clark et al., 2005, Ferreira et al., 2023].

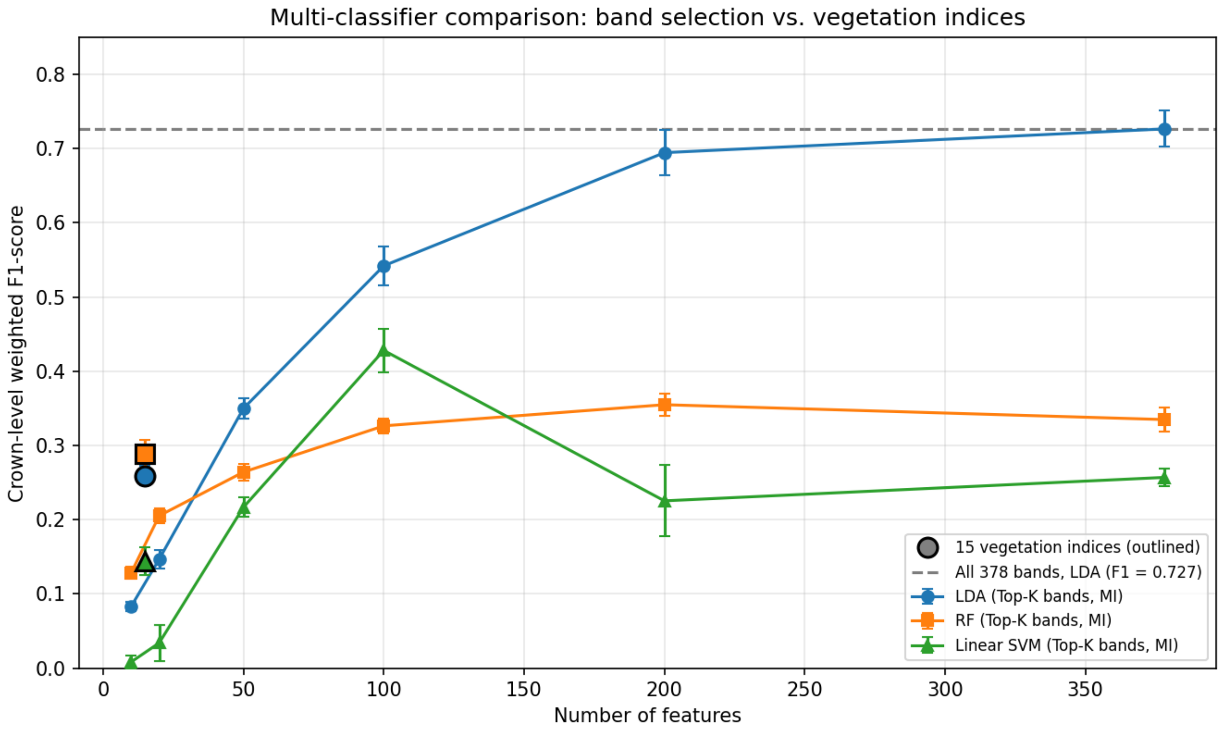


**Figure S8:** Multi-classifier comparison across model-agnostic feature sets. Lines show LDA (blue), Random Forest (orange), and linear SVM (green) trained on MI-ranked Top-$K$ bands. Outlined markers show performance on 15 standard vegetation indices. LDA dominates at all feature counts, with the performance gap widening as dimensionality increases. Dashed line: full-band LDA baseline (F1 = 0.727).

#### S7 Illumination threshold sensitivity

The study's preprocessing pipeline removed pixels with illumination fraction below 60%, following Schläpfer et al. [2015]. To test whether this threshold materially affects classification performance, we re-ran the full LDA pipeline (stratified-group 5-fold cross-validation with crown-level majority vote, matching Section 2.7) at five illumination thresholds: 0% (no filtering), 20%, 40%, 60% (default), and 80%.

Weighted F1-score was remarkably stable across the entire range (Fig. S9; Table S4), varying by only 0.6 percentage points between the best (60%, F1 = 0.727) and worst (20%, F1 = 0.721) thresholds. Macro F1-score showed a similarly flat pattern. Even with no illumination filtering (0%), accuracy (weighted F1 = 0.726) was within the error bars of the default threshold, despite including 22% more pixels. At 80%, three species lost their minimum crown count of two and were excluded, yet accuracy remained comparable (weighted F1 = 0.721). These results indicate that classification accuracy is robust to the choice of illumination threshold across a wide range, and that the spectral signal distinguishing species is not strongly confounded by within-crown shadow variation at this spatial resolution.

**Table S4:** Classification accuracy across illumination thresholds. Weighted and macro F1 are means $\pm$ SD over 5 folds. Note: this sensitivity sweep used a rasterised 178-species pixel snapshot, which includes 9 species absent from the canonical train/test split owing to a processing-pipeline version difference. All headline results in the main text and the canonical CV (Section 2.7, Fig. 8) use the 169-species set.

| Threshold (%) | Pixels | Crowns | Species | Weighted F1 |
|---|---|---|---|---|
| 0 | 387,206 | 2,781 | 178 | 0.726 $\pm$ 0.022 |
| 20 | 341,862 | 2,780 | 178 | 0.721 $\pm$ 0.010 |
| 40 | 331,311 | 2,780 | 178 | 0.724 $\pm$ 0.015 |
| 60 | 318,546 | 2,780 | 178 | 0.727 $\pm$ 0.024 |
| 80 | 294,800 | 2,729 | 175 | 0.721 $\pm$ 0.012 |

#### S8 Band importance by taxonomic family

To test whether the most important spectral bands for classification differ across major taxonomic groups – addressing the concern that "optimal bands may not be suitable for the classification of all tree species" – we trained separate LDA models for each of the six most species-rich families (families with $\geq$ 3 species each having $\geq$ 10 crowns: Fabaceae, Sapotaceae, Vochysiaceae, Lecythidaceae, Clusiaceae, and Anacardiaceae) and extracted per-band importance using the variance-weighted squared scalings method described in Section S6.1.

The resulting importance profiles (Fig. S10) show that all six families concentrate discriminative information in the same far-red-edge region ($\sim$730–780 nm) that dominates the global (all-species) importance profile. However, families differ in the sharpness and exact location of their importance peaks. Sapotaceae shows the most concentrated peak (centred at $\sim$760 nm), while Fabaceae and Lecythidaceae spread importance more broadly into the VIS and NIR, reflecting greater functional diversity among species within those families. Anacardiaceae (the smallest family with only 3 species) shows a narrower and shifted peak.

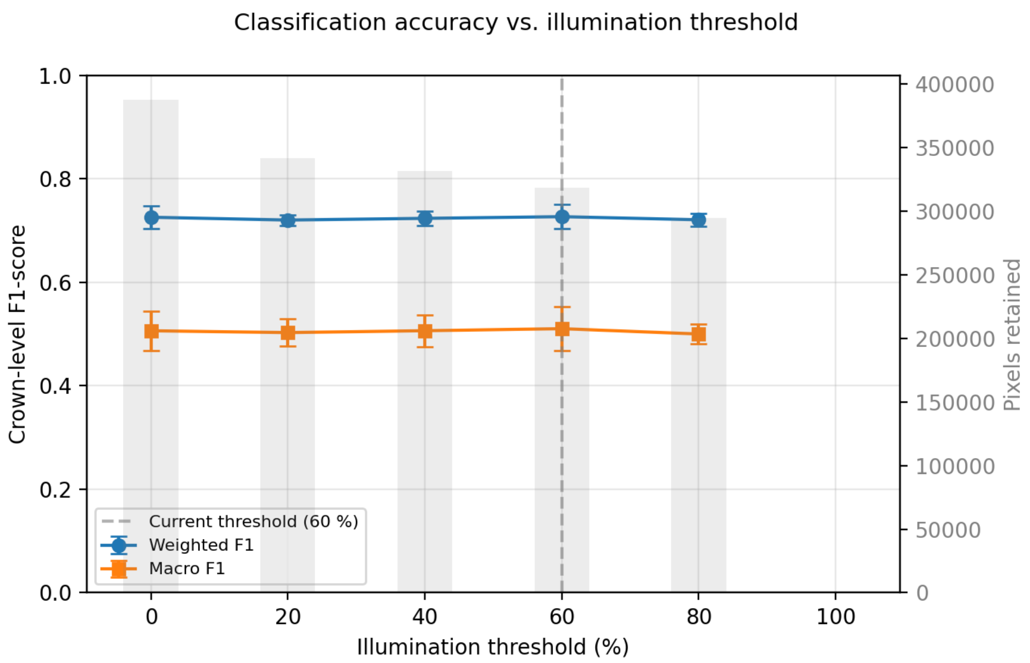


**Figure S9:** Crown-level classification accuracy as a function of the illumination filtering threshold. Blue circles: crown-level weighted F1-score (mean ± SD over 5 folds). Orange squares: macro F1. The dashed vertical line marks the default 60% threshold used in this study. Grey bars show the number of pixels retained at each threshold. Accuracy is stable across the full 0–80% range, with only 0.6 pp variation in weighted F1.

These results indicate that while the FRE is universally the most informative spectral region for within-family species discrimination at this site, the relative importance of secondary spectral features varies by lineage. This is consistent with the observation that functional trait variation is structured by phylogeny [Ball et al., 2026] and provides partial support for the hypothesis that different taxonomic groups rely on distinct biochemical markers for spectral differentiation.

#### S9 Confusion matrix on held-out test set

To visualise which species are confused by the LDA classifier on the held-out test set (Section 2.7), we computed the row-normalised crown-level confusion matrix using `caret::confusionMatrix` (Fig. S11). Entries are predicted-class fractions per true-class row, so cells along the diagonal denote per-species recall and off-diagonal cells reveal systematic misclassification patterns. Species are grouped along both axes by taxonomic family (coloured side strips), with families ordered by descending species richness. The dominant pattern is a strong diagonal for common species and increasing off-diagonal mass for species with few training crowns; most off-diagonal mass falls within or between closely related families, consistent with the phylogenetic conservatism of canopy spectra discussed in Section S5.2.

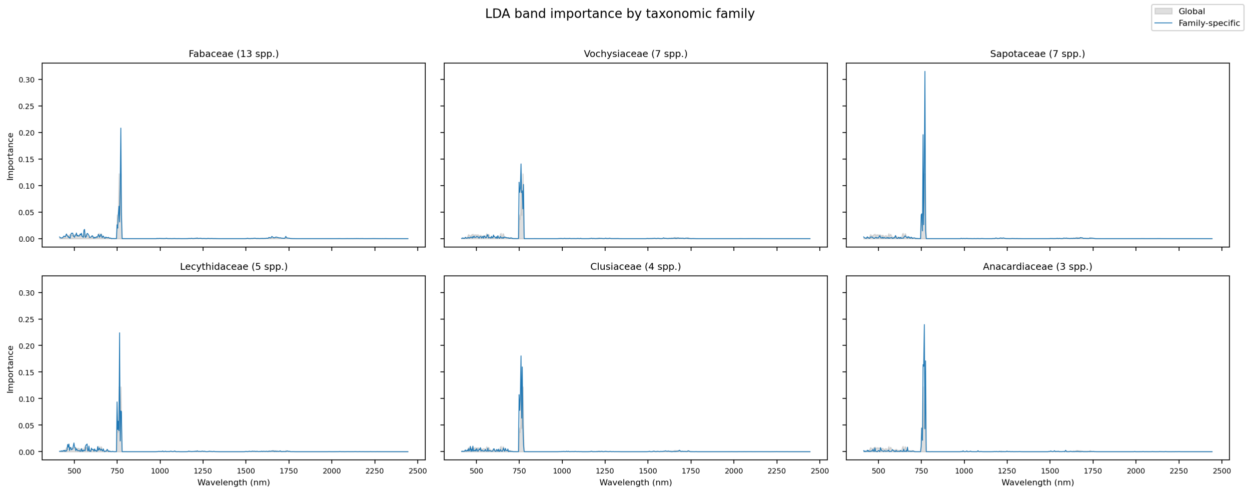


**Figure S10:** LDA band importance for the six largest taxonomic families compared to the global (all-species) model. Each panel shows the variance-weighted squared scalings for a family-specific LDA (blue line) overlaid on the global importance profile (grey fill). All families concentrate discriminative power in the far-red-edge (∼730–780 nm), but differ in peak sharpness and secondary features. Number of species per family (with $\geq$ 10 crowns) shown in panel titles.

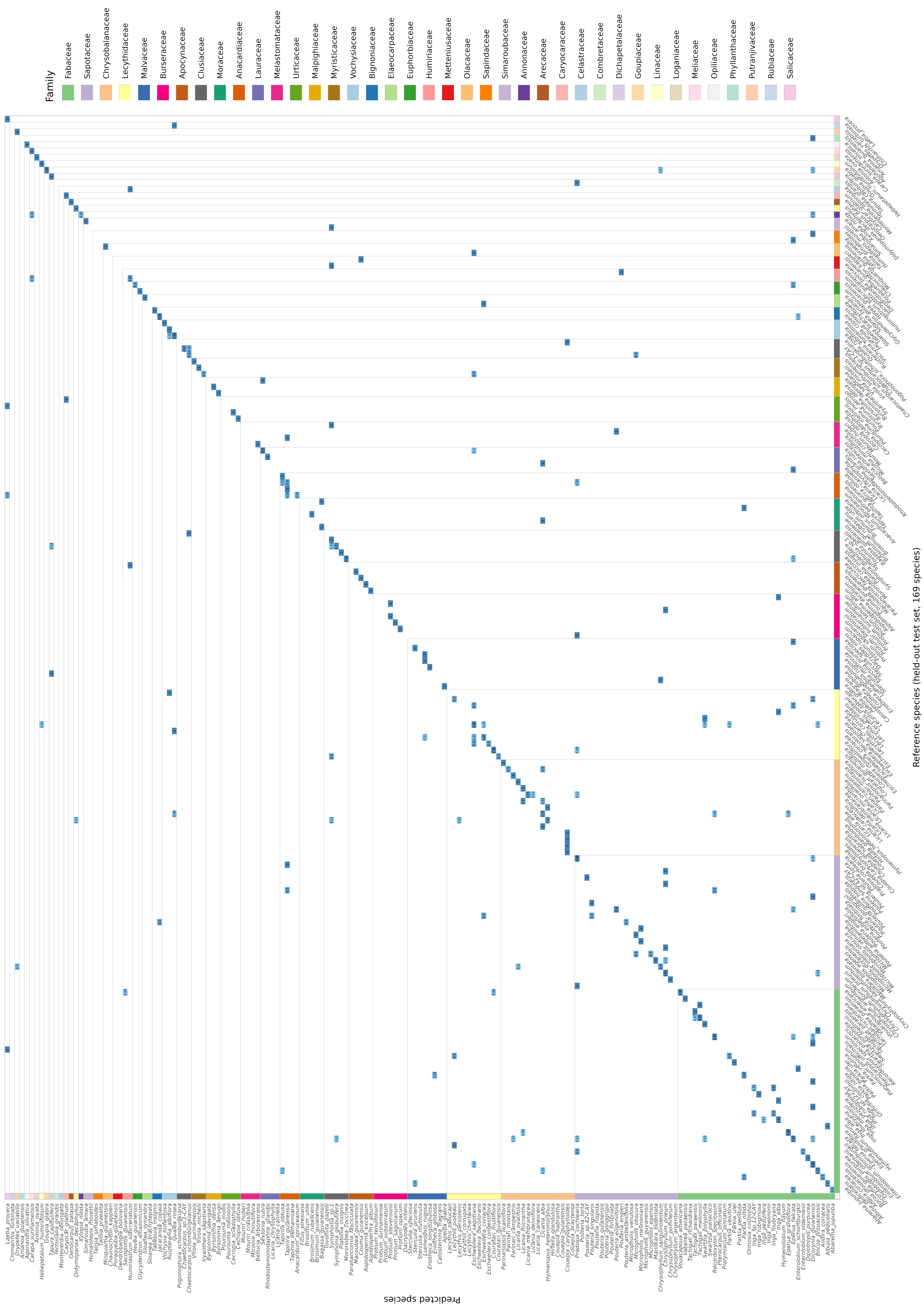


**Figure S11:** Row-normalised crown-level confusion matrix for the LDA classifier on the held-out test set (169 species, 666 test crowns; overall crown-level accuracy 0.78). Each row sums to 1; cell colour indicates the fraction of true-class crowns predicted as each species. Species are ordered along both axes by taxonomic family (coloured side strips, families ordered by descending species richness, 41 families in total). Most off-diagonal mass concentrates within (or between adjacent) families, reflecting the phylogenetic conservatism of canopy spectra.
