## Supplementary figures and images for "Advances and limits of species-level canopy mapping in a hyperdiverse tropical forest: multi-temporal crown segmentation and airborne imaging spectroscopy"

### Supplementary Confusion Matrix

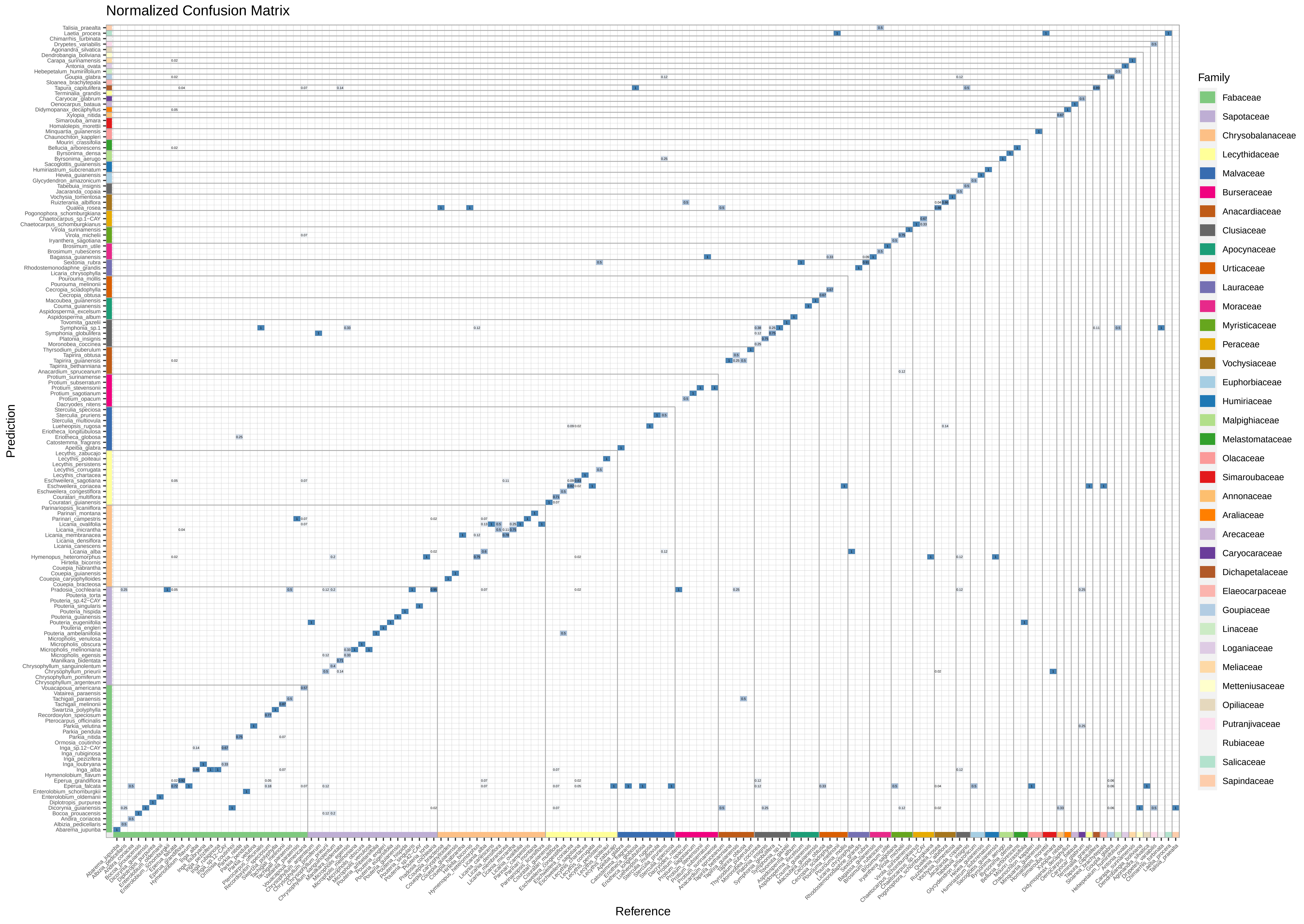
